# Glycolysis downregulation is a hallmark of HIV-1 latency and sensitizes infected cells to oxidative stress

**DOI:** 10.1101/2020.12.30.424810

**Authors:** Iart Luca Shytaj, Francesco Andrea Procopio, Mohammad Tarek, Irene Carlon-Andres, Hsin-Yao Tang, Aaron R. Goldman, MohamedHusen Munshi, Mattia Forcato, Konstantin Leskov, Fengchun Ye, Bojana Lucic, Nicolly Cruz, Amit Singh, Silvio Bicciato, Sergi Padilla-Parra, Marina Lusic, Ricardo Sobhie Diaz, David Alvarez-Carbonell, Andrea Savarino

**Author notes:** Equal contribution (David Alvarez-Carbonell, Andrea Savarino).

## Abstract

HIV-1 infects lymphoid and myeloid cells, which can harbor a latent proviral reservoir responsible for maintaining lifelong infection. Glycolytic metabolism has been identified as a determinant of susceptibility to HIV-1 infection, but its role in the development and maintenance of HIV-1 latency has not been elucidated. By combining transcriptomic, proteomic and metabolomic analysis, we here show that transition to latent HIV-1 infection downregulates glycolysis, while viral reactivation by conventional stimuli reverts this effect. Decreased glycolytic output in latently infected cells is associated with downregulation of NAD^+^/NADH. Consequently, infected cells rely on the parallel pentose phosphate pathway and its main product, the antioxidant NADPH, fueling antioxidant pathways maintaining HIV-1 latency. Of note, blocking NADPH downstream effectors, thioredoxin and glutathione, favors HIV-1 reactivation from latency in lymphoid and myeloid cellular models. This provides a “shock and kill effect” decreasing proviral DNA in cells from people-living-with-HIV/AIDS. Overall, our data show that downmodulation of glycolysis is a metabolic signature of HIV-1 latency that can be exploited to target latently infected cells with eradication strategies.

## Introduction

Decades after its outbreak, the HIV/AIDS pandemic remains one of the main causes of morbidity and mortality of humankind, leading to almost one million victims per year (source: UNAIDS). Moreover, due to the severe medical and economic crisis caused by Coronavirus Disease 2019 (COVID-19), adherence to and availability of antiretroviral therapies (ART) is decreasing in areas where HIV/AIDS prevalence is particularly high (Jiang *et al*, 2020; Jewell *et al*, 2020). This worsening of the death toll highlights the fragility of current therapeutic approaches, based on lifelong ART administration. Therefore, a cure for HIV/AIDS is an unmet medical need of growing importance for people-living-with-HIV/AIDS (PLWH). The quest for a cure is hampered by persistence of transcriptionally silent proviruses within latently infected cells, which render these cells hard to discriminate from their uninfected counterparts (Finzi *et al*, 1999). Latent HIV-1 DNA can be mainly found in viral reservoirs such as CD4^+^ T-cells (Van Lint *et al*, 2013), however myeloid cells (in particular microglia) can also contribute to persistence of the infection during ART (Sattentau & Stevenson, 2016). Pinpointing molecular features to allow selective targeting of latently HIV-1 infected cells would represent a significant, and perhaps decisive, step in the quest for an HIV/AIDS cure.

A possible approach to reach this goal is the investigation of the metabolic pathways exploited by the retrovirus to actively replicate and enter a latent state. In this regard, it is interesting to point out that the cells more susceptible to HIV-1 infection are characterized by an increased glycolytic rate (Valle-Casuso *et al*, 2019). Cellular activation, leading to increased glycolysis, is necessary for active HIV-1 replication; of note, in CD4^+^ T-cells and monocytes of HIV-1 infected individuals, glucose consumption is increased and the expression of the glucose transporter GLUT-1 is upregulated (Palmer *et al*, 2014a, 2014b).

Conversely, reports on the effect of HIV-1 infection on glycolysis have been sparse and, in some respects, conflicting. Starting from the 1990s, studies have shown that cells infected with HIV-1 display decreased levels of NAD^+^, an important substrate of glycolytic reactions (Murray *et al*, 1995), and that supplementation of the NAD^+^ precursor nicotinamide influences viability of productively HIV-1 infected cells (Savarino *et al*, 1996). Moreover, after ART implementation, PLWH showed to be more susceptible to hyperglycemia than the general population (Dubé *et al*, 1997). On the other hand, upregulated glucose metabolism was reported to favor apoptosis of infected CD4^+^ T-cells (Hegedus *et al*, 2014). Although these combined sets of data strongly support a role of active glycolysis in determining susceptibility to HIV-1 infection as well as an altered glucose metabolism in PLWH, the role of glycolysis in viral latency, and therefore in possible curing strategies, has as yet remained unclear.

The importance of understanding the relationship between cell metabolism and the infection is emphasized by the possible first long-term remission of HIV-1 without bone marrow transplantation (Abstract Supplement Oral Abstracts from the 23rd International AIDS Conference 2020). This individual had been treated with a combination of intensified ART and the NAD^+^ precursor nicotinamide, thus suggesting a possible contribution of glycolysis regulation in the therapeutic result obtained. The anecdotal character of this case, however, renders difficult the drawing of definitive conclusions.

Indirect evidence of an interplay between glycolysis and HIV-1 latency can also be drawn by the dysregulation of redox pathways, which are intertwined with glycolytic metabolism in several cell types and pathological conditions (Locasale & Cantley, 2011; Kondoh *et al*, 2007). This interconnection might be relevant, as we recently showed that HIV-1 infection leads to enhancement of antioxidant defenses in primary CD4^+^ T-cells (Shytaj *et al*, 2020). HIV-1 infection causes an initial oxidative stress (Daussy *et al*, 2020), which then leads to the nuclear translocation of the master antioxidant transcription factor Nrf2. This translocation in turn induces transcription of several proteins involved in antioxidant response, including glucose 6-phosphate dehydrogenase (G6PDH), which diverts glucose 6-phosphate from the glycolytic pathway to the pentose cycle, responsible for production of the antioxidant NADPH. Therefore, elucidating the specific glycolysis/redox state interconnection during HIV-1 infection would be of pivotal importance for understanding the molecular events which lead infected cells to either die or establish a latent infection.

Herein, we combine transcriptomic, proteomic, and metabolomic datasets, including single cell analyses, to show that, during transition to HIV-1 latency, glycolysis is downregulated. In line with this, our results show that latently infected cells able to undergo HIV-1 reactivation are also characterized by prompter reactivation of glycolysis. Moreover, we show that downregulation of glycolysis in latently infected cells is accompanied by higher reliance on the antioxidant thioredoxin (Trx) and glutathione (GSH) systems for cell survival. Our results highlight the possibility to exploit glycolytic imbalances induced by HIV-1 infection for the elimination of retrovirally infected cells. This result may improve our knowledge of pathways that can be targeted by strategies aimed at eradicating HIV/AIDS.

### HIV-1 infection downregulates expression of glycolytic enzymes in CD4^+^ T-cells

To study transcriptomic profiles upon infection, we used microarray and RNA-Seq data sets generated in primary CD4^+^ T-cells infected with HIV-1_pNL4-3_. The cellular model employed is based on longitudinal sample collection to cover different time points, from early HIV-1 infection [day 3 post-infection (p.i.)] to peak retroviral replication (days 7-9 p.i.) and latency/survival of infected cells (day 14 p.i.). The detailed features and validations of this cellular model have been described previously elsewhere (Shytaj et al, 2020).

Microarray results highlighted significant downregulation of the glycolytic pathway in infected cell cultures, which was independently evidenced by Gene Set Enrichment Analysis [GSEA; (Subramanian et al, 2005)] either using a customized gene set comprising enzymes involved in human glycolysis (henceforth, HUMAN-GLYCOLYSIS) or using the Reactome or Biocarta databases (Figure 1A; Additional file 1). Entrance of the infected cells into a hypometabolic state was also shown by downregulation of glycolysis-independent transcriptional and translational pathways, although downregulation of these pathways did not reach the same level of convergence among the databases examined as compared to the glycolytic pathway (Additional file 1). Moreover, among the pathways more heavily perturbed by HIV-1 infection, there was the interferon pathway (Additional file 1), as expected, and pathways associated with both apoptosis and cell cycle, in line with the fact that, in an HIV-1 infected cell cultures, some cells succumb to infection and others survive developing a proviral latent state.

**Figure 1.**
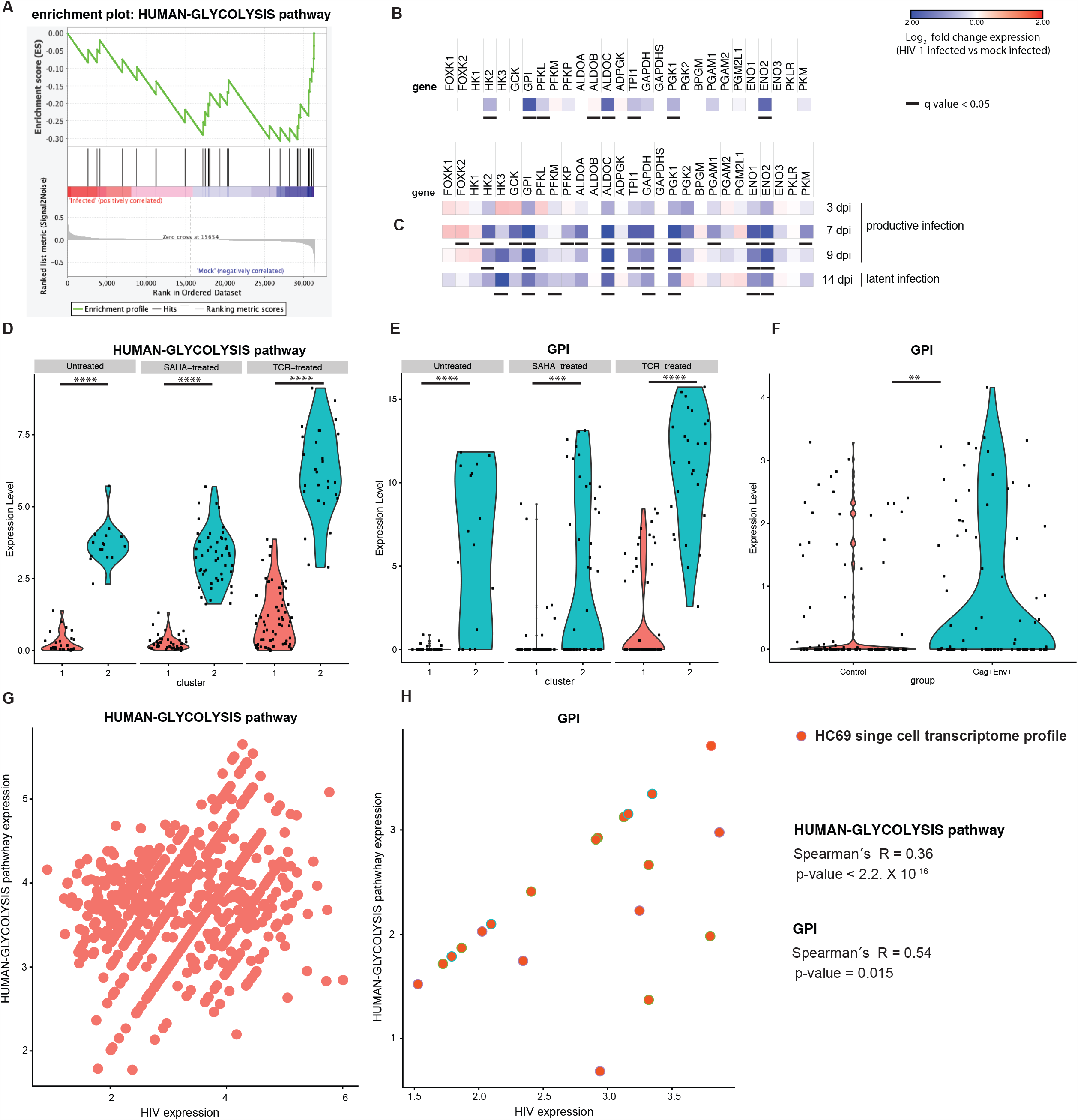
Transcriptional modulation of glycolytic enzymes in productively or latently HIV-1 infected cells. Panels A-C) primary CD4^+^ T-cells were activated with α-CD3-CD28 beads and infected with HIV-1_pNL4-3_ or mock-infected. Cells were cultured for two weeks post-infection (p.i.) to model different infection stages (days 3-9 p.i., *i*.*e*. productive infection; day 14 p.i., *i*.*e*. latent infection) and subjected to microarray (A,B; n=2) or RNA-Seq (C; n=3) analysis. Panel A) Gene set enrichment analysis (GSEA) of the expression of the glycolytic pathway (HUMAN-GLYCOLYSIS) in mock-infected or HIV-1 infected cells. B,C) Heatmaps of the relative expression of glycolytic enzymes upon HIV-1 infection. Data are expressed as Log_2_ fold change in HIV-1 infected vs mock-infected cells. For microarray data (B) expression values of infected and mock-infected cells at different time points were pooled. For RNA-Seq data (C), expression values in infected cells were normalized using the corresponding time point in mock-infected cells. Adjusted p values (q values) to account for multiple testing were calculated by Significance Analysis of Microarrays [SAM (Tusher *et al*, 2001)] and Deseq2 for RNA-Seq data (Love *et al*, 2014). Panels D-F) scRNA-Seq of the expression of the entire glycolytic pathway or of glucose phosphate isomerase only (*GPI*) in primary CD4^+^ T-cells infected *in vitro* (D,E) or CD4^+^ T-cells of PLWH (F). In panels D,E, cells were infected with VSVG-HIV-1-GFP and sorted for viral expression as detailed in (Golumbeanu *et al*, 2018). Following latency establishment, cells were left untreated or HIV-1 expression was reactivated through suberoyl anilide hydroxamic acid (SAHA) or α-CD3-CD28 engagement. Clusters 1 and 2 were identified by principal component analysis as described in (Golumbeanu *et al*, 2018). In panel F, CD4^+^ T-cells were isolated from total blood of PLWH under ART as described in (Cohn *et al*, 2018). Viral expression was reactivated by treatment with phytohemagglutinin (PHA) and cells were sorted using antibodies against Env and Gag. Sorted cells were then subjected to scRNA-Seq analysis. The expression level of the HUMAN-GLYCOLYSIS pathway in Panel D was calculated as the average expression of genes comprising the gene list; expression levels in cluster 1 and 2 were compared using Wilcoxon rank sum test. For panels E-F significance of *GPI* differential expression level between clusters (E) or between control and Env^+^ Gag^+^ conditions (F) was assessed by Wilcoxon rank sum test encoded in FindMarkers Seurat R function. Panels G,H) Correlation of combined HIV-1 expression and the expression of the entire glycolytic pathway (G) or *GPI* only (H) in sc-RNA-Seq profiling of untreated HC69 microglial cells. Data were analyzed by Spearman’s correlation coefficients. ** p< 0.01, *** p< 0.001; *** p< 0.0001.

We then analyzed in detail the glycolytic enzymes of the pathway HUMAN-GLYCOLYSIS (Figure 1B). The glycolytic enzymes characterized by significantly downregulated transcription upon infection were hexokinase 2 (*HK2*), glucose 6-phosphate isomerase (*GPI*), phosphofructokinase liver type (*PFKL*), aldolase fructose-bisphosphate C (*ALDOC*), triosephosphate isomerase 1 (*TPI1*) and enolase 2 (*ENO2*), thus suggesting a broad downregulation of the glycolytic pathway (Figure 1B). In particular, the highly significant downregulation of *GPI* suggests a glycolysis-specific effect, as this enzyme commits metabolites to the glycolytic, rather than to the alternative pentose-phosphate pathway.

To expand these analyses, we explored an RNA-Seq dataset derived from the same primary CD4^+^ T-cell model, which was previously published elsewhere (Shytaj et al, 2020). In this dataset, the number of donors and time points was higher, allowing the study of the expression of glycolytic enzymes during each infection stage (Figure 1C). In line with the microarray results, differential gene expression analysis of RNA-Seq data (DESeq2) (Love et al, 2014) highlighted significant downregulation of glycolysis in infected cells cultures, which was initiated at peak HIV-1 replication (7-9 days p.i.) and persisted after retroviral replication had ceased (14 days p.i.) (Figure 1C). As seen in microarray analysis, enzyme transcriptional downregulation covered all main steps of glycolysis, and the RNA-Seq data set further suggested that various isoforms, in particular of *PFK*, might contribute differently to glycolysis downregulation during productive or latent infection [*PFK*-platelet (P); adjusted p value = 0.04 at 7 days p.i. *PFK*-muscle (M); adjusted p value = 0.04 at 14 days p.i. respectively]. Finally, analysis of a previously published proteomic dataset of the same CD4^+^ T-cell model (Shytaj et al, 2020) further corroborated the downregulating effect of HIV-1 on glycolysis, confirming significant downmodulation of GPI, PGK1 and TPI1 (Additional file 2).

Overall, these data show that HIV-1 infection during its transition to latency is associated with downregulated expression of glycolytic enzymes, in particular those catalyzing the early steps of glycolysis.

### Expression of glycolytic enzymes is required for HIV-1 escape from latency in lymphoid and myeloid cells

The aforementioned results prove that glycolysis downregulation is initiated during productive infection and accompanies HIV-1 latency establishment. We then proceeded to specifically investigate the transcriptional regulation of the HUMAN-GLYCOLYSIS pathway upon the reverse process, *i*.*e*. reactivation from latency.

To this aim, we first analyzed two single cell RNA-Seq (scRNA-Seq) datasets independently published by the groups of Ciuffi and Nussenzweig, respectively (Golumbeanu *et al*, 2018; Cohn *et al*, 2018). In the first dataset, primary CD4^+^ T-cells infected with pseudotyped HIV-1-GFP/VSVG had been sorted according to GFP expression and allowed to revert to latency (Golumbeanu *et al*, 2018). Latently infected cells had then been either left untreated or subjected to HIV-1 reactivation by strong (α-CD3/CD28 antibodies) or weak stimuli [suberoyl anilide hydroxamic acid (SAHA)]. Eventually, the transcriptomic profile was analyzed by scRNA-Seq (Golumbeanu *et al*, 2018). Using principal component analysis, the authors identified two cell clusters, which were less (cluster 1) or more (cluster 2) susceptible to HIV-1 reactivation (Golumbeanu *et al*, 2018). We analyzed the expression of the enzymes of the glycolytic pathway in both clusters and found that glycolytic enzymes were downregulated in cells of cluster 1 (Figure 1D). Interestingly, this relative downregulation, already visible in basal conditions, was maintained upon treatment with either anti-CD3/CD28 antibodies or SAHA (Figure 1D). Moreover, *GPI* expression was lower in the cell subpopulation less responsive to HIV-1 reactivation as compared to the more susceptible cell subpopulation (Figure 1E), in line with the results obtained on our model of latency establishment (Figure 1B,C).

To further validate the clinical relevance of these findings, we analyzed a second scRNA-Seq dataset obtained from CD4^+^ T-cells of PLWH under ART. To generate this dataset, the Nussenzweig group separated cells responsive to HIV-1 reactivation by detecting Gag/Env expression upon treatment with the pan-lymphocyte activator, phytohemagglutinin (PHA) (Cohn *et al*, 2018). Despite its limitations (the lower number of infected cells as compared to the previously mentioned model and the likely presence of a mixed population of infected and uninfected cells), this dataset had the advantage of capturing a viral reservoir generated under the pathophysiologic conditions occurring *in vivo* in a clinical setting. In line with the results shown by sc-RNA-Seq of *in-vitro* infected cells, the cell population responsive to HIV-1 reactivation displayed significant upregulation of *GPI* (Figure 1F).

We then expanded our analysis to include microglial cells transformed with SV40 and latently infected with HIV-1 [*i*.*e*. HC69 cells (Garcia-Mesa *et al*, 2017; Alvarez-Carbonell *et al*, 2017)]. Microglia constitutes one main myeloid retroviral reservoir during ART and is largely responsible for HIV-1 persistence in the central nervous system (Churchill & Nath, 2013). Our bulk RNA-Seq HC69 data, along with data on its uninfected equivalent (C20 cells) allowed a comparison of reactivated *vs*. latent infection through stimulation of latently infected cells with tumor necrosis factor [TNF (Garcia-Mesa *et al*, 2017)]. When infected cells were compared to their uninfected counterparts incubated under similar conditions, results showed a clear pattern of *GPI* downregulation being more significant in latent than in productive infection and accompanied by a more pronounced downregulation of the early glycolytic enzymes in the former (Additional File 3). Finally, sc-RNA-Seq profiling of HC69 cells highlighted a significantly positive correlation between baseline HIV-1 expression and key glycolytic enzymes, including *GPI*, in the subset of cells in which proviral transcription was detectable (Figure 1G,H). Conversely, treatment with dexamethasone (DEXA), which is a known glycolysis inhibitor (Ma *et al*, 2013), decreased both the proportion of HC69 cells expressing the transcripts of the glycolytic pathway and the baseline percentage of cells expressing HIV-1 transcripts (Additional file 4).

Taken together, these results show that low expression of early glycolytic enzymes, in particular *GPI*, is associated with HIV-1 latency maintenance, and that at least partial restoration of glycolysis is required for latency disruption.

### Decreased glycolytic metabolism during productive and latent HIV-1 infection

To connect transcriptional downregulation of glycolytic enzymes with metabolic profiling, we first used a recently developed FRET-based biosensor system in living TZM-bl cells infected with HIV-1 (San Martín *et al*, 2013). We combined this reporter system with labelled HIV-1 Gag to measure the stability of the glycolysis surrogate marker, lactate, at a single cell resolution using Fluorescence Lifetime Microscopy (FLIM) (Coomer *et al*, 2020) (Figure 2A). Data showed a significant decrease of lactate levels in cells infected with HIV-1JR-FL 3 dpi, as compared to No Env infected cells. This decrease was similar to the reduction in lactate levels observed upon glycolysis inhibition by 2-deoxy-D-glucose (2-DG) treatment (Figure 2B), corroborating the finding that downregulation of the glycolysis pathway is accompanied by decreased output of glycolytic metabolites following HIV-1 infection.

**Figure 2.**
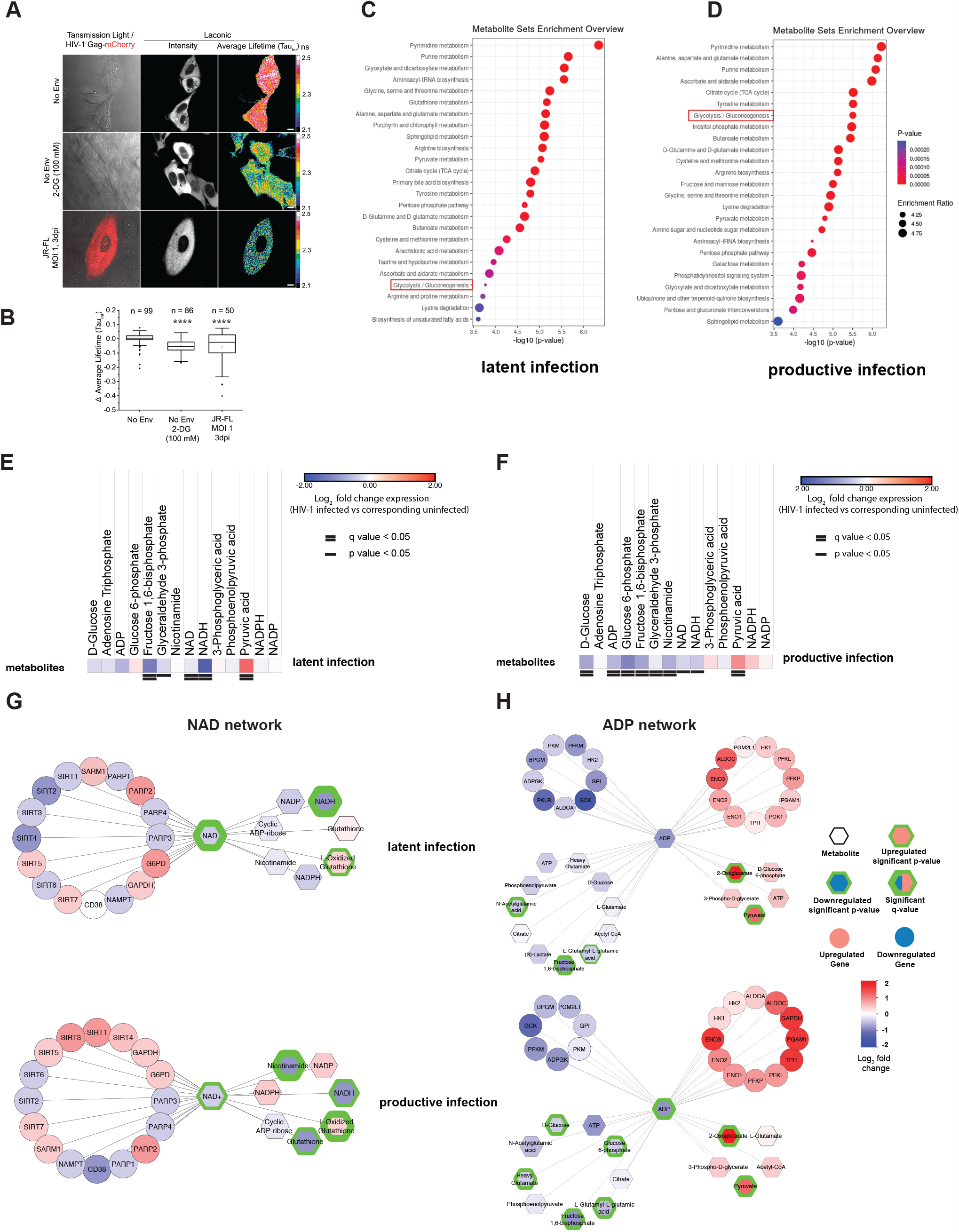
Modulation of glycolytic metabolites during productive or latent HIV-1 infection. Panel A) micrograph showing TZM-bl cells expressing Laconic FRET-based biosensor infected with HIV-1 Gag-mCherry pseudoviruses (No Env or JR-FL Env, as indicated), in the absence or presence of the glycolysis inhibitor D-deoxy D-glucose (2-DG). Left column: merge of transmission light image and Gag-mCherry fluorescence showing TZM-bl cells negative (-mCherry) or positive for HIV-1 infection (+mCherry). Middle column: intensity image of Laconic fluorescence. Right column: pixel-by-pixel average lifetime (TauInt) image of Laconic expressing cells represented in 16-colors Look-up Table (LUT). Calibration color bar shows values from 2.1 to 2.5 ns. Scale bar = 10 µm. Panel B) Box and whisker plot showing results from individual TZM-bl cells from 3 independent experiments. Total number of cells analyzed per condition is indicated above each box. Results were analyzed using one-way ANOVA and Sidak post-hoc test ****: p < 0.0001. Panels C,D) Metabolite enrichment analysis in latently (C) and productively (D) infected microglial cells as compared to their uninfected counterparts. The top enriched pathways were ordered according to p values obtained with Q statistic for metabolic datasets performed with Globaltest (MetaboAnalyst) (Xia *et al*, 2009). Panels E,F) Heatmap of glycolytic metabolites in HC69 latently infected cells (E) or HC69-TNF cells with HIV-1 reactivated (F) as compared to their uninfected counterparts. Data are displayed as Log_2_ fold change expression. P values were calculated by Student’s t-test while the Benjamini-Hochberg false discovery rate was used to account for multiple testing and calculate q values. Panels G,H) Network analysis of metabolites and genes associated with NAD^+^/NADH (G) and ADP (H) pathways. Genes differentially expressed are derived from the RNA-Seq analysis shown in Additional file 3. Hexagons represent metabolites, circles represent genes and the figure legend describes the differentiation pattern of nodes. Networks were built using Cytoscape software (http://www.cytoscape.org) using Metscape plugin (http://metscape.ncibi.org/tryplugin.html) (Gao *et al*, 2010).

To discern the metabolic profiles of productive and latent HIV-1 infection, we then subjected the aforementioned microglial model to LC-MS/MS metabolomic profiling (Figure 2C-G; Additional files 5,6). The choice of this model allowed investigating a homogeneously infected cell population which, nevertheless, preserves many features of primary microglia (Garcia-Mesa *et al*, 2017). As mentioned above, four conditions could be analyzed: quiescent, activated, latently infected, and productively infected.

Metabolite enrichment analysis showed significant changes in the glycolysis/gluconeogenesis pathway in both latently and productively infected cells as compared to their uninfected counterparts (Figure 2C,D). We then proceeded to analyze the glycolytic metabolites separately.

Results from latently infected cells showed significantly decreased levels of the metabolite produced by PFK, *i*.*e*. fructose 1,6 bisphosphate (Figure 2E). Since the PFK enzyme acts immediately downstream of GPI, downregulation of fructose 1,6 bisphosphate is in line with the reduced expression of *GPI* in latently infected cells, as shown by both RNA-Seq and scRNA-Seq analysis (Figure 1). There was also a trend towards reduction of glyceraldehyde 3-phosphate, a product of ALDO, the enzyme acting immediately downstream of PFK (Figure 2E). These results, again, support the view that latently infected cells are characterized by a block in the early glycolytic steps.

Instead, the pattern observed in cells reactivated from latency, as compared to uninfected (C20) cells subjected to the same HIV-1 reactivating stimulus, showed a less clear impairment of glucose metabolism, with parameters indicative of glycolysis downregulation (a relative paucity of the initial glycolytic metabolites) and parameters indicative of glucose consumption (significantly decreased D-glucose levels and decreased ADP levels) (Figure 2F).

We then explored the connection between decreased glycolytic metabolism and other, intertwined, metabolic pathways highlighted by the aforementioned enrichment analysis (Figure 2C,D and 2G,H; and Additional files 5,6). Apart from the purine/pyrimidine pathways (likely related to DNA synthesis and retrovirus/host interactions), productively and latently infected cells shared increased glutamate metabolism, among the top-10 significantly enriched pathways (Figure 2C,D; Additional files 5,6). Network analysis supports the view that these cells rely on glutamate metabolism as an alternative source of pyruvate for fueling ADP utilization through the Krebs cycle (Figure 2H).

Among the most significantly enriched pathways, we further considered metabolism of antioxidants (Additional file 5), due to its well described interconnection with glycolysis and NAD^+^, via the pentose pathway (Grant, 2008), and its relevance in the establishment and maintenance of HIV-1 latency (Shytaj *et al*, 2020; Benhar *et al*, 2016). Network analysis revealed a likely connection between NADH decrease and redox pathways in latently infected cells (Figure 2G). Accordingly, increased levels of oxidized glutathione, consistent with higher oxidative stress upon infection, as previously described (Shytaj *et al*, 2020), were associated with a decrease not only in NADH but also in its precursor NAD^+^. The HIV-1-related NAD^+^ consumption was also associated with altered nicotinamide metabolism (Additional files 5,6) and with a trend towards increased expression of proteins consuming NAD^+^/NADH (Figure 2G). Among the most up- or downregulated transcripts (> 1 or < −1 Log_2_ fold) involved in this pathway were *G6PD* and *PARP2* in latent infection, the former being the enzyme initiating the pentose phosphate pathway and the latter being a NAD^+^-consuming enzyme involved in repair of oxidative-stress induced DNA damage (Chevanne *et al*, 2007) (Figure 2G). Moreover, productively, but not latently, infected cells were characterized by higher levels of sirtuin (*SIRT*)1 and 3, which are known to favor HIV-1 replication (Pagans *et al*, 2005) and to counteract mitochondrial oxidative stress, respectively (Singh *et al*, 2018) (Figure 2G). Productively infected cells also displayed decreased expression of *CD38*, an ectoenzyme responsible for hydrolysis of extracellular NAD^+^ and intracellular recycling of its components, ADP-ribose and nicotinamide (Savarino *et al*, 2000) (Figure 2G).

Overall, these data show that early glycolytic metabolites are downregulated in both latently and productively HIV-1 infected cells, as compared to uninfected cells. Moreover, the results indicate that HIV-1 infection diverts metabolism from glycolysis to the pentose cycle, as further described below.

### Downstream block of the glycolysis-alternative pentose phosphate pathway can induce a “shock and kill” effect in latently infected cells

The data so far presented suggest that glycolysis is downregulated in latently HIV-1 infected cells to divert glucose 6-phosphate to the pentose cycle. In this regard, the standard balance between the two pathways would be altered by NAD^+^ paucity (Figures 2E-G), as NAD^+^ is not only a main cofactor for glycolytic enzymes, but also a precursor of the main pentose cycle cofactor NADP^+^ (Xiao *et al*, 2018). As diversion to the pentose phosphate pathway leads to regeneration of the antioxidant NADPH, we tested whether blocking the downstream effectors of NADPH could induce HIV-1 escape from latency and exploit NADH paucity to reduce the ability of infected cells to survive oxidative stress.

For this purpose, we chose the drugs auranofin (AF) and buthionine sulfoximine (BSO), which by inhibiting thioredoxin (Trx) and glutathione (GSH) regeneration, respectively (Benhar et al. 2016), can block the antioxidant/pro-latency effect of NADPH. These drugs were also preferred because of their translational potential, due to their clinical (separately) and pre-clinical (combined) testing as anti-reservoir compounds in PLWH and macaques infected with the simian immunodeficiency virus (SIV) (Diaz *et al*, 2019; Benhar *et al*, 2016). As expected, at the concentrations chosen for the reactivation experiments, the two drugs were able to synergistically increase oxidative stress in a previously described reporter model which allows measuring GSH potential in live cells (Bhaskar *et al*, 2015) (Additional File 7). When we analyzed HIV-1 production in a number of proviral latency models, we found that AF/BSO favored proviral reactivation, at different efficiencies, in both lymphoid and myeloid models (Figure 3A, Additional Files 8,9).

**Figure 3.**
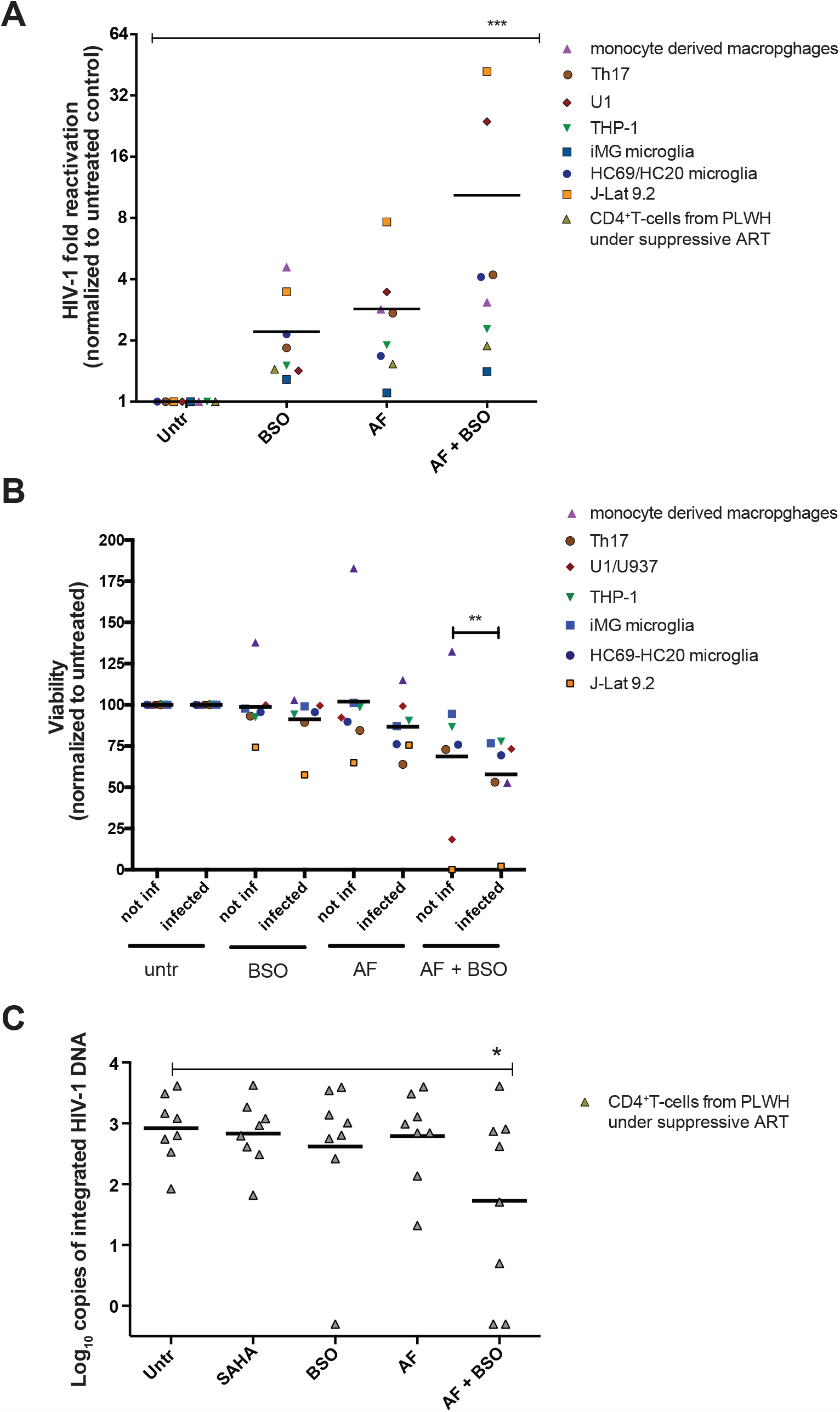
“Shock and kill” effect of combined thioredoxin (Trx) and glutathione (GSH) inhibition in cellular models of HIV-1 latency and CD4^+^ T-cells of PLWH. Panels A,B. Reactivation from HIV-1 latency (A) and relative cell viability (B) in different cell models following treatment with the Trx inhibitor auranofin (AF; 500 nM), the GSH inhibitor buthionine sulfoximine (BSO; 250 μM), or a combination of the two. The characteristics of the different models adopted are detailed in the Material and Methods section. Each data point represents a mean from at least two independent experiments conducted in the different models. Replicates of all experiments for each cell model are shown in Additional Files 8, 9 and 11, except for the data of monocyte-derived macrophages which were retrieved from (Shytaj *et al*, 2013). Panel C. Levels of integrated HIV-1 DNA following treatment for 48 h with AF and/or BSO in CD4^+^ T-cells derived from PLWH under suppressive antiretroviral therapy. Live cells were sorted after treatment, and integrated DNA was measured by Alu-PCR. The latency reactivating agent SAHA was used as a reference compound (Archin *et al*, 2012). Data were analyzed by non-parametric Friedman’s test followed by Dunn’s post-test (A,C) or two-way ANOVA followed by Tukey’s post-test (B). Solid lines represent the means. * p< 0.05, ** p< 0.01, *** p< 0.001.

Moreover, when cell viability was analyzed, results showed that combined inhibition of Trx and GSH led to the preferential killing of HIV-1 infected cells as compared to their uninfected counterparts (Figure 3B), although specific leukemia/lymphoma cell lines were highly sensitive to AF and BSO treatment irrespective of HIV-1 infection, in line with the previously described sensitivity of such neoplasias to these drugs (Benhar *et al*, 2016; Fiskus *et al*, 2014).

We finally tested the combination of AF and BSO in primary cells derived from PLWH under ART. Analysis of metabolic profiles of this primary cell model showed significant enrichment of the pentose cycle upon treatment with both AF and BSO as compared to cells treated with BSO only (Additional file 10). These effects were not detectable when cells treated with AF-only were compared to cells treated with BSO, thus supporting the specificity of the drug combination. Both results are in line with the well-known compensation of GSH inhibition by the Trx system (Benhar *et al*, 2016), thus confirming that only the combination of both drugs can lead to a sustained pro-oxidant effect. Finally, we tested the therapeutic potential of AF and BSO on CD4^+^ T-cells of PLWH receiving ART. As, in these *ex-vivo* samples, direct and precise analysis of cell viability (infected vs. control) was not possible due to the low frequency of HIV-1-infected cells, we indirectly tested the selective survival/elimination of HIV-1 infected cells by measuring the frequency of the integrated proviral DNA in cells sorted for viability after treatment. Results showed that, despite only minor effects on HIV-1 reactivation as measured by *Tat/rev* Induced Limiting Dilution Assay [TILDA (Procopio *et al*, 2015)] (Additional file 11), proviral HIV-1 DNA was significantly lower in cell cultures that had received both AF and BSO (Figure 3C). Of note, cells from two of the donors showed loss of integrated proviral DNA signal after AF/BSO treatment.

Overall, these data show that glycolysis downregulation and increased reliance on the pentose cycle in latently HIV-1 infected cells can be exploited by pro-oxidant drugs targeting Trx and GSH to induce HIV-1 reactivation and/or mortality of infected cells.

## Discussion

The results of the present study identify downregulation of glycolytic activity as a determinant of the transition from productive to latent HIV-1 infection, an effect that is reversed upon proviral reactivation. Our data show that it is HIV-1 infection *per se* that paves the way towards glycolysis downregulation, since productively HIV-1-infected cells display glycolytic profiles intermediate between those of uninfected and latently infected cells. Of note, HIV-1-associated downregulation of glycolysis was shown to occur mainly due to downmodulation of glycolysis-initiating enzymes (*e*.*g*. GPI) and paucity of precursors of the energetic products of glycolysis.

Our findings complement previous studies showing that the cell subsets most susceptible to HIV-1 infection are those displaying higher glycolytic activity and that this metabolic state supports HIV-1 replication (Valle-Casuso *et al*, 2019; Hegedus *et al*, 2014). The present study was instead aimed at a longer follow up, comprising transition to proviral latency from productive infection and proviral reactivation from latency. The overall evidence points toward a model where progressive downregulation of glycolysis is associated with gradual dampening of retroviral replication, perhaps selecting those cells that can survive the productive phase and undergo latent infection. The fact that latently infected cells undergo a metabolic transition, is further corroborated by our results showing that conventional HIV-1 reactivation stimuli can preferentially enhance glycolysis upon reversion to productive infection (Figure 4).

**Figure 4:**
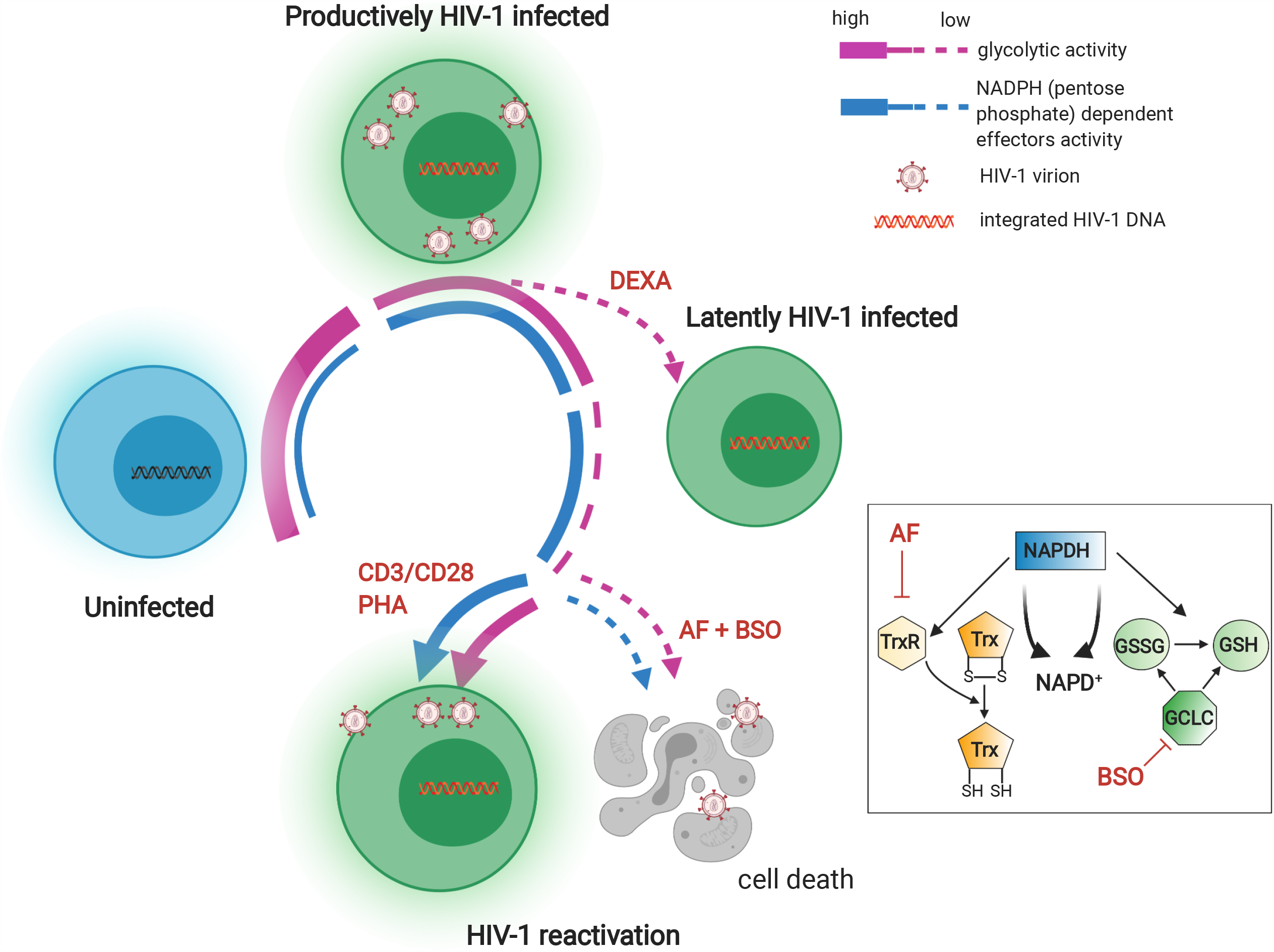
Glycolysis downregulation is a hallmark of HIV-1 latency and sensitizes infected cells to oxidative stress. Schematic model of the regulation of glycolysis during sequential stages of HIV-1 infection and of its potential therapeutic targeting (created with BioRender). Uninfected cells become susceptible to HIV-1 infection upon activation by upregulating a number of metabolic pathways including glycolysis (Valle-Casuso *et al*, 2019). During productive HIV-1 infection, glycolysis is downregulated, a process which is in part compensated by enhancement of the parallel pentose phosphate pathway. Latent HIV-1 infection can ensue either spontaneously, leading to further glycolysis downregulation, or be induced by agents such as dexamethasone (DEXA), which can directly inhibit glycolysis. During latency, the pentose phosphate pathway remains active and fuels, through NADPH activity, regeneration of the two main antioxidant molecules, *i*.*e*. Trx and GSH. Reversion to a productively infected state is accompanied by partial reactivation of the glycolytic pathway. Blocking Trx and GSH inhibits downstream effects of NADPH without restoring glycolytic activity, thus favoring death of latently infected cells. TrxR = thioredoxin reductase; GCLC = Glutamate-cysteine ligase; GSSG = oxidized GSH.

One study that reached a partially different conclusion from ours is the work of Castellano *et al*. which analyzed a time course of macrophages infected *in vitro* with HIV-1 (Castellano *et al*, 2019). This study concluded that glycolysis might not be impaired in latently infected macrophages, although it pinpointed that these cells rely on glycolysis-independent energy sources such as glutamate. The authors based their conclusions on the analysis of the acidification capability (a byproduct of glycolysis). Our report, however, shows decreased production of lactate at the single cell level, which may have been missed using a cell culture containing HIV-1 infected cells surrounded by a majority of uninfected bystander cells. On the other hand, our report confirms that, when glycolysis is downregulated, glutamate becomes a readily alternative source for pyruvate in latently infected cells, so as to maintain sufficient levels of OXPHOS-derived ATP production compatible with cell survival. In this regard, another study analyzed a panel of 186 different metabolites in plasma of PLWH and highlighted glutamate was one out of the 6 metabolites that was unequivocally elevated among PLWH as compared to uninfected control individuals (Scarpelini *et al*, 2016).

Our analysis of the glycolytic pathway at a single cell resolution may reconcile our conclusions with previous results obtained studying the metabolism of PLWH. In this regard, Palmer *et al*. showed that glycolysis is upregulated in PLWH, independently of ART (Palmer *et al*, 2014a, 2014b). As the percentage of infected cells *in vivo* is relatively low, this effect could be attributed to chronic immune hyperactivation (Deeks *et al*, 2004), rather than to retroviral replication *per se*. In line with a possible contribution of bystander effects to the *in-vivo* observations, the data sets analyzed in the present work clearly showed transcriptional downregulation of the glycolysis pathway in single cells characterized by a deep latency state.

A hallmark of HIV-1 infection identified by our metabolomic analysis is the consumption of nicotinamide, which is a NAD^+^ precursor. This molecule is also necessary for synthesis of NADP^+^, the precursor of the antioxidant NADPH, which is the main pentose cycle product. Therefore, nicotinamide consumption may contribute to divert metabolism from glycolysis to the pentose cycle, consistent with a model where glycolysis favors HIV-1 replication while the pentose cycle favors HIV-1 latency as well as antioxidant defenses. The latter phenomenon is in line with the previously described upregulation of antioxidant responses in latently HIV-1 infected cells (Shytaj *et al*, 2020). Moreover, oxidative stress immediately following HIV-1 infection was shown to induce increased transcription of G6PDH (Shytaj et al. 2020), therefore further validating the diversion of glucose 6-phosphate to the pentose cycle. Moreover, these observations may shed new light on the previously reported contribution of nicotinamide to proviral escape from latency (Samer *et al*, 2020) and viability of HIV-1-infected cells (Savarino *et al*, 1997).

Our data also show that reinforcing glycolysis downmodulation through pro-oxidant drugs acting downstream of NADPH may lead to the selective elimination of infected cells. In particular, NADPH acts as a cofactor for the regeneration of the two most important antioxidant defenses, Trx and GSH (Benhar *et al*, 2016; Miller *et al*, 2018). In line with this, dual inhibition of the Trx and GSH pathways might induce a vicious cycle: on one side, subtracting metabolites to glycolysis and readdressing them to the antioxidant defense-generating pentose cycle, and on the other side blocking the downstream antioxidant effects of the pentose cycle (Figure 4). In this regard, an interesting parallel can be drawn with neoplastic cells, as previous studies showed increased susceptibility to the cytotoxic effects of combined Trx and GSH pathway inhibition in lung cancer cells with pharmacologically inhibited glycolysis (Fath *et al*, 2011; Li *et al*, 2015). Indeed, while our study privileged the use of a dual drug combination to decrease the likelihood of non-specific toxicity, it is tempting to speculate that including an inhibitor of glycolysis could enhance the effects observed on latently infected cells. Interestingly, the use of one such inhibitor (2DG) alone was able to decrease viral replication in the study of Valle Casuso *et al*. in line with our model where downregulation of glycolysis favors the latent/non productive phase of the infection (Valle-Casuso *et al*, 2019).

A possible limitation of our study is that it focused mainly on glycolysis as the energy-producing pathway. In this regard, another major source of NADH is the Krebs cycle (Wu *et al*, 2015; Viña *et al*, 2016), which intervenes downstream of glycolysis. We believe that the complexity of these phenomena will require stepwise analysis. Future work might shed light on the regulation of the Krebs cycle during latent HIV-1 infection and thus provide a more complete and integrated picture of the metabolic alterations induced by HIV-1 infection.

The translational relevance of our findings resides in the fact that the drugs selected to inhibit the Trx and GSH pathways could become a useful tool for inducing a “shock-and-kill” effect, helping to purge the latently infected cells (Deeks, 2012). Indeed, the drugs used in the present work (AF and BSO) had already shown the potential to decrease the viral reservoir in macaques and PLWH (Shytaj *et al*, 2015; Diaz *et al*, 2019). A similar effect was observed in macaques treated with another prooxidant compound and Trx pathway inhibitor, *i*.*e*. arsenic trioxide (Yang *et al*, 2019). One characteristic of these strategies was not only the elimination of latently infected cells *in vivo* but also the enhancement of anti-HIV-1 cell mediated immunity (Shytaj *et al*, 2015). In light of the results of the present study, this immune enhancement could be interpreted as a consequence of the “shock” effect against viral latency provided by the combined inhibition of the Trx and GSH pathways. Future studies will be required to test this hypothesis and to understand the potential interplay between oxidative stress and cell-mediated immunity in the context of HIV-1 infection. Overall these results highlight glycolysis downregulation as a distinctive metabolic feature of latently HIV-1 infected cells which can be exploited to target latent reservoirs.

## Materials and Methods

### Cell cultures and HIV-1 infection

The following cell lines or primary cells were used as models of productive or latent HIV-1 infection: 1) lymphoid cells: a) latently HIV-1 infected J-Lat 9.2 cells and uninfected Jurkat T-cells, b) latently HIV-1 infected primary Th17 cells (Alvarez-Carbonell *et al*, 2017), c) primary CD4^+^ T-cells from healthy donors infected with HIV-1 *in vitro* (Shytaj *et al*, 2020) and d) primary CD4^+^ T-cells from PLWH with viral loads stably suppressed by ART; 2) myeloid cells: a) the THP-1 monocytic cell line, b) the U937 and U1 (latently HIV-1 infected) promonocytic cell lines, c) primary human monocyte derived macrophages (Shytaj *et al*, 2013), d) human immortalized microglia (hµglia) clone C20 (uninfected) and clone HC69 (latently infected with HIV-1) (Garcia-Mesa *et al*, 2017; Llewellyn *et al*, 2018; Alvarez-Carbonell *et al*, 2017), e) iPSC-derived microglia (iMG) uninfected or infected with HIV-1 (iMG/HIV) (Alvarez-Carbonell *et al*, 2019); 3) reporter TZM-bl cells transfected with HIV-1 Gag-mCherry viruses pseudotyped with JR-FL Env or without Env.

#### 1. Lymphoid cells

The parental Jurkat E6.1 (ATCC number: TIB-152) cell line and the latently HIV-1 infected J□Lat 9.2 cell line (Jordan et al, 2003) were cultured using RPMI + 10% fetal bovine serum (FBS) and plated at 0.25–0.5 × 10^6^ cells/mL as previously described (Shytaj *et al*, 2020).

Latently HIV-1 infected Th17 cells were generated as previously described (Dobrowolski *et al*, 2019; Garcia-Mesa *et al*, 2017). Briefly, naive CD4^+^ T cells were isolated using a RoboSep CD4^+^ Naïve T cell negative selection kit (STEMCELL Technologies Inc., Vancouver, British Columbia, Canada), and 2□X□10^6^ cells were resuspended in 10□mL RPMI medium and stimulated with 10□µg/mL concanavalin A (ConA) (EMD Millipore, Billerica, MA, USA) in the presence of subset-specific cytokines. Cells were cultured for 72□h at 37°C, followed by addition of 10□mL of fresh medium, additional 10□µg/mL ConA, polarization cocktail cytokines, and 120□IU/mL of IL-2. After 6□days, cells were washed and resuspended in RPMI medium supplemented with the growth cytokines IL-23 (50□ng/mL) and IL-2 (60□IU/mL). Cells were then infected in a 24-well plate using VSV glycoprotein-pseudotyped virus expressing CD8 and GFP at a multiplicity of infection (MOI) of 2 at a cellular concentration of 5□X□10^6^ cells per mL, in the presence of cell subset cytokines. Cells were spinoculated at 2,000□×□*g* for 1.5□h at room temperature and then placed in an incubator overnight. Cells were adjusted to 1□×□10^6^ per mL in the presence of cell subset-appropriate growth cytokines. After 48□h, infection efficiency was determined by GFP expression and infected cells were isolated using RoboSep mouse CD8a positive selection kit II (STEMCELL Technologies Inc., Vancouver, British Columbia, Canada). Cells (50□X□10^6^ per mL) were pre-incubated with 50□µL/mL of antibody cocktail from the kit and 40□µL/mL of magnetic beads and diluted into 2.5□mL RoboSep buffer. Positive cells were recovered by magnetic bead separation, suspended in 1□mL of medium, and vortexed to release the cells and beads from the tube wall.

Primary CD4^+^ T-cells for *in-vitro* HIV-1 infection were isolated from total blood of healthy individuals using the RosetteSep™ Human CD4^+^ T Cell Enrichment Cocktail (STEMCELL Technologies Inc., Vancouver, British Columbia, Canada) as previously described (Shytaj *et al*, 2020) and according to the manufacturer’s instructions. The blood was obtained through the Heidelberg University Hospital Blood Bank following approval by the local ethics committee. To induce activation before HIV-1 infection Dynabeads® Human T□Activator CD3/CD28 was added to cells for 72 h. Cells were then mock-infected or infected using 2ng p24 of HIV□1_pNL4_ 3/10^6^ cells. Mock-infected and HIV-1 infected cells were then cultured for two weeks in RPMI + 20% FBS with 10 ng/mL IL□2 at a density of 0.5-2 × 10^6^/mL. At 3,7,9 and 14 days post-infection 1 × 10^6^ cells were pelleted and used for RNA extraction and transcriptomic analysis (microarray and RNA-Seq) as previously described (Shytaj *et al*, 2020).

For *ex-vivo* experiments on CD4^+^ T cells or PBMCs of PLWH, cells from adult donors were analyzed. The use of samples was approved by the Institutional Review Board of the Centre Hospitalier Universitaire Vaudois and by the Human Subjects Review Committee of the Federal University of Sao Paulo. All subjects gave written informed consent. Experiments were exclusively performed on cells isolated from treated PLWH with undetectable viremia (HIV-1 RNA levels <50 copies per mL of plasma) for at least 12 months. Culture conditions and HIV-1 reactivation experiments were conducted as described in (Procopio *et al*, 2015).

#### 2. Myeloid cells

For producing latently HIV-1 infected THP-1 (ATCC number: TIB-202) cell cultures, uninfected cells were cultured on a 6-well plate at a density of 1 × 10^6^ cells per well in RPMI growth medium containing 10% FBS, 1% penicillin/streptomycin, and 50 nM of 2-mercaptoethanol. Infection with a HIV-1-GFP virus was carried out by spinoculation, as described in (Alvarez-Carbonell *et al*, 2017). Positively selected cells were placed in RPMI medium with cell type-specific growth cytokines at 1□X□10^6^ in upright flasks and allowed to expand for a week prior to treatments. During all assays suspension cells were cultured at a density of 1 × 10^6^ cells per mL, in 96-well plates in a volume of 100 µL.

Monocytic cell lines U937 and U1 (harbouring the latent HIV-1 provirus) were cultured in RPMI + 10% FBS. Cells were stably transduced to express the Grx1-roGFP2 biosensors in cytoplasm as described previously (Bhaskar *et al*, 2015).

The procedure used to generate hµglia/HIV HC69 from C20 was previously described in (Garcia-Mesa *et al*, 2017). Cells were cultured in BrainPhysTM medium (STEMCELL Technologies Inc., Vancouver, British Columbia, Canada) containing 1X N2 supplement-A (Gibco-Invitrogen, #17502–048), 1X penicillin streptomycin (Gibco ™, #15140122), 100 μg/mL normocin™ (Invivogen, #ant-nr-1), 25 mM glutamine (Gibco ™, #25030081), 1% FBS, and 1 μM DEXA (freshly added to the cell culture) (Sigma-Aldrich, # D4902), as previously described (Garcia-Mesa *et al*, 2017). For experiments cells were plated at 0.1 × 10^6^ cells per well in 24-well plates.

Human iPSC-derived microglia (Tempo-iMG™; Tempo Bioscience, CA) was cultured in DMEM/F-12 (Thermo Fisher, Waltham, MA, USA) supplemented with 1X N2 supplement, 0.5X NEAA, 2 mM L-Glutamine, 100 ng/mL GM-CSF, 50 ng/mL IL-34, and infected with VSVG-HIV-GFP as previously described (Alvarez-Carbonell *et al*, 2019). The experiments with iMG and iMG/HIV cells (adherent) were performed 72h post-infection at a density of 50,000 cells per well in a 96-well plate pre-coated with Growth Factor Reduced Matrigel (Corning Inc., Corning, NY, USA).

#### 3. TZM-bl cells

TZM-bl (kindly provided by Dr. Quentin Sattentau, University of Oxford) and Lenti-XTM 293T cells (Takara Bio, Clontech, Saint Germain en Laye, France) were cultured in either complete DMEM or complete DMEM F-12 (Thermo Fisher, Waltham, MA, USA), respectively, supplemented with 10% FBS and 1% penicillin-streptomycin. Cells were grown at 37°C in the presence of 5% CO_2_.

For HIV-1 pseudovirus production, the pR8ΔEnv plasmid (encoding HIV-1 genome harbouring a deletion within *env*), pcRev and NL4-3 Gag-mCherry ΔEnv were kindly provided by Dr. Greg Melikyan (Emory University, Atlanta, GA, USA). The plasmid encoding JR-FL Env was a kind gift from Dr James Binley (Torrey Pines Institute for Molecular Studies, FL, USA). For lactate measurements, Laconic construct was obtained from Addgene (ref. 44238).

HIV-1 pseudovirus bearing Gag-mCherry was produced as described previously (Coomer *et al*, 2020). Briefly, Lenti-XTM 293T cells were transfected at 60–70% confluency with a mix of pR8ΔEnv, pcRev, NL4-3 Gag-mCherry ΔEnv and with or without JR-FL Env, at a 2:1:3:3 ratio, using GeneJuice transfection reagent (Novagen, EMD Millipore, Billerica, MA, USA). 72 h after transfection, viral supernatants were collected, filtered (0.45□μm) and concentrated using Lenti-XTM Concentrator (Takara Bio, Clontech, Saint Germain en Laye, France).

Tzm-bl were plated onto 8-well µ-Slide (Cat.No: 81826, Ibidi, Gräfelfing, Germany) and transfected on the same day with 250 ng of plasmid expressing the Laconic biosensor per well, using GeneJuice transfection reagent. 12 h post-transfection, the medium was replaced with fresh complete DMEM after washing with PBS 1X and cells were incubated for further 4 h at 37°C. 16 h post-transfection, cells were infected at MOI of 1 with HIV-1 Gag-mCherry viruses pseudotyped with JR-FL Env or without Env as a negative control for infection (No Env viruses). Viruses were diluted in a final volume of 100 μL FluoroBrite DMEM 2% FBS (Thermo Fisher, Waltham, MA, USA) per well and added onto cells. Cells were spinoculated at 2,100 *g* in a refrigerated centrifuge (4°C) for 20 min. The viral inoculum was then removed, cells were washed with PBS 1X and incubated for further 90 min in FluoroBrite DMEM 2% FBS at 37°C to allow viral fusion to occur. The medium was later replaced by complete DMEM and cells were again incubated at 37°C. Transfected cells expressing Laconic, challenged to HIV-1 pseudoviruses, were analyzed by Fluorescence Lifetime Microscopy (FLIM) 3 days post-infection as described below.

### RNA extraction

Total cellular RNA was extracted using the InviTrap® Spin Universal RNA Mini Kit (Stratec Biomedical, Germany) according to the manufacturers’ instructions and as previously described (Shytaj *et al*, 2020). RNA concentration was assessed using a P-class P 300 NanoPhotometer (Implen GmbH, Munich, Germany).

### Microarray and RNA-Seq analyses

Primary CD4^++^T-cells infected with HIV-1 or mock-infected were subjected to microarray and RNA-Seq using 500 ng of total RNA that was quality checked for integrity via Bioanalyzer. Microarray was performed using the HumanHT-12 beadchip (Illumina, Inc., 5200 Illumina Way San Diego, CA 92122 USA) and scanned using an iScan array scanner. Data extraction was done for all beads individually, and outliers were removed when the absolute difference to the median was greater than 2.5 times MAD (2.5 Hampelís method). Raw data are available at GSE163405.

Bead-level microarray raw data were converted to expression values using the lumi R package for quality control, variance stabilization, normalization, and gene annotation (Du *et al*, 2008). Briefly, raw data and control probes were loaded in R using the lumiR and addControlData2lumi functions. Raw signals were background corrected, estimating the background based on the control probe information with the bgAdjust method of the lumiB function. Background corrected data were then processed with the variance-stabilizing transformation (VST) of the lumiT function to stabilize the variance and were finally normalized using the quantile normalization implemented in lumiN. Gene expression data were annotated using the R package illuminaHumanv4.db that contains the mappings between Illumina identifiers and gene descriptions.

To identify the impact of HIV-1 infection on gene expression, we compared the expression levels of CD4^+^ T-cells infected with HIV-1 with those of mock-infected cells using Significance Analysis of Microarray (Tusher *et al*, 2001) algorithm coded in the same R package. In SAM, we estimated the percentage of false-positive predictions (*i*.*e*., false discovery rate, FDR) with 100 permutations and selected as differentially expressed those genes with an FDR q-value ≤ 0.05.

Over-representation analysis was performed using Gene Set Enrichment Analysis (Subramanian *et al*, 2005) and using the gene sets of the Biocarta and Reactome collections from the Broad Institute Molecular Signatures Database (http://software.broadinstitute.org/gsea/msigdb) as well as a customized gene set derived from AmiGO (see section “Glycolysis pathway” below). GSEA software (http://www.broadinstitute.org/gsea/index.jsp) was applied on Log_2_ expression data of cells infected with HIV-1 or matched mock-infected controls. Gene sets were considered significantly enriched at FDR < 5% when using Signal2Noise as metric and 1,000 permutations of gene sets.

Differential expression of genes in RNA-Seq datasets was analyzed using the DESeq2 package (Love *et al*, 2014). The datasets used were retrieved from (Shytaj *et al*, 2020) (<u>GSE127468</u>) for CD4^+^ T-cells and from (Garcia-Mesa *et al*, 2017) (<u>SRP075430</u>) for uninfected (C20) and HIV-1 infected (HC69) microglia. Heatmaps were generated using the Morpheus tool (https://software.broadinstitute.org/morpheus).

### Sc-RNA-Seq analyses

scRNA-Seq datasets were retrieved from (Golumbeanu *et al*, 2018) (<u>GSE111727</u>) and from (Cohn *et al*, 2018) (<u>GSE104490</u>) and analyzed with the Seurat [version 3.1.5; (Stuart *et al*, 2019)] R package. In particular, normalized gene expression data for GSE111727 were downloaded from Zenodo repository (Zenodo_Data_S2) and loaded into a Seurat object; raw gene counts for GSE104490 were downloaded from GEO and normalized using Seurat. Differentially expressed genes were calculated using the FindMarkers function of the Seurat package.

The scRNA-Seq data of HC69 (GSE163979) were generated as follows: immortalized human microglia HC69 cells were left untreated or treated with dexamethasone [DEXA, 1μM (Sigma-Aldrich, St. Louis, MO, USA)] for 72 h. After harvesting, a minimum of 600,000 viable cells per condition was subjected to Drop-Seq as described in (Macosko *et al*, 2015). After capturing individual cells in the oil droplets with barcoded beads (Chemgenes, Inc. Wilmington, MA, 01887 USA), droplets were broken, and cDNA libraries were generated using Illumina Nextera XT kit (Illumina, Inc., 5200 Illumina Way San Diego, CA, 92122 USA). Next Generation Sequencing and quality control were performed at Medgenome Inc (Foster City, CA, 94404 USA). Drop-Seq Tools v.1.0 and STAR-2.5.1b alignment tools were used to process and map the sequences to the hg18 human reference genome and to the retroviral portion of the HIV-1_pNL4-3_ construct. As a result, we built digital gene expression matrices (DGE) containing the read counts for human and HIV-1 genes. The DGE matrices were used to generate the Seurat objects. After filtering, a total of 6,528 control and 5,869 DEXA-treated individual cell transcriptome profiles were consolidated, each expressing at least 100 genes, with each gene expressed in at least 3 cells. To prevent “zero inflation” bias, further filtering was performed to isolate only cells expressing HIV-1 and the genes of the glycolytic pathway (HUMAN-GLYCOLYSIS). Correlation scatter plots were built using Seurat, accompanied by the calculations of the Spearman correlation coefficients R and related p-values.

### Proteomic analysis

Proteomic analysis of CD4^+^ T-cells infected *in vitro* with HIV-1 or mock-infected was retrieved from (Shytaj *et al*, 2020). The dataset is available at the ProteomeXchange Consortium via the PRIDE (Perez□Riverol et al, 2019) partner repository with the dataset identifier PXD012907 (http://www.ebi.ac.uk/pride/archive/projects/PXD012907). Relative protein quantification was performed by Despite Data-independent Acquisition (DIA) processing raw data with Spectronaut Pulsar X (version 11) using default and previously described parameters (Shytaj *et al*, 2020). Heatmaps were generated using the Morpheus tool (https://software.broadinstitute.org/morpheus).

### Metabolomic analysis

Metabolomic analysis was performed as described previously (Li *et al*, 2020). Briefly, polar metabolites were extracted from cells using a cold extraction solution containing 80% methanol and 20% water. Two types of analyses were conducted. One set was run undiluted (1x), while the other set was concentrated 5-fold before analysis (5x). To increase sensitivity of detection, the 5x dataset was employed, except for heat sensitive metabolites of interest (*i*.*e*. glutathione and D-glucose), which were analyzed starting from the undiluted run. Samples were analyzed by LC-MS/MS on a Thermo Q Exactive HF-X mass spectrometer coupled to a Vanquish LC System. LC separation was performed by hydrophilic interaction liquid chromatography (HILIC), pH 9, using a SeQuant ZIC-pHILIC column (MilliporeSigma). Peak areas, representing metabolite levels, were extracted using Thermo Compound Discoverer 3.0. Peak areas were normalized to protein amounts calculated on the lysates. Metabolites were identified by accurate mass and retention based on pure standards and by accurate mass and MS/MS fragmentation followed by searching the mzCloud database (www.mzcloud.org). For heatmap generation, Log_2_ fold change values were calculated based on average expression values from triplicate samples. Statistically significant changes were assessed by Student’s *t*-test (p-value) and the Benjamini-Hochberg false discovery rate (FDR) to account for multiple testing (q-value).

Metabolite pathway and enrichment analysis were performed using MetaboAnalyst 4.0 (http://www.metaboanalyst.ca) (Xia *et al*, 2009). For enrichment analysis, a table of raw peak intensity values was submitted to the platform and the Benjamini-Hochberg FDR correction was used to adjust all p-values and reduce false-positive discovery for multiple testing. The q conversion algorithm was then used to calculate FDRs in multiple comparisons using FDR < 0.05 as threshold for significance in all tests.

For pathway analysis, Human Metabolome Database (HMBD) (Wishart, 2020) IDs of the metabolites differentially expressed between the conditions compared were submitted to the MetaboAnalyst platform. Comparisons were analyzed by Student’s t-test using Holm’s correction for multiple comparisons, so as to generate a q-value. The library ‘Homo sapiens (human)’ of the Human Metabolome Database was used for pathway analysis. For network generation, metabolite data were integrated with the RNA-Seq data of the same microglia model (described in its dedicated methods section). As input data, KEGG IDs, p-values and Log_2_ fold changes were used for the selected compounds (metabolites and gene transcripts). In order to analyze a correlation network of the compounds in shared pathways, MetScape (Gao *et al*, 2010), an app implemented in Java and integrated with Cytoscape (version 3.5.1), was used.

### Glycolysis pathway

Except for *a priori* analysis (GSEA and metabolic pathway enrichment), the list of enzymes and metabolites used to define the glycolytic pathway was selected based on known literature and to reflect all metabolic/enzymatic steps associated with glycolysis irrespective of the cell type examined. Specifically, the gene list was retrieved from AmiGO Gene ontology using the filters: GO:0061621 (canonical glycolysis) AND Homo sapiens (pathway named in the paper as: HUMAN-GLYCOLYSIS). The metabolite list was retrieved from the KEGG Pathway database entry M00001 [Glycolysis (Embden-Meyerhof pathway)].

### Drug treatments

For glycolysis inhibition, TZM-bl cells infected with JR-FL without Env (No Env virus) were treated with 100 mM 2-deoxy-glucose (2-DG) (Sigma-Aldrich, St. Louis, MO, USA) 2 h prior to image acquisition.

For viral reactivation and cell viability experiments, cells were treated with auranofin (Sigma Aldrich #A6733; 500 nM), L-Buthionine-sulfoximine (*i*.*e*. Sigma Aldric BSO #B2515; 250 μM), or a combination of the two for 24 or 4 8h as indicated in the captions of Additional files 8,9 and 11. In parallel, cells were incubated with one the following positive control activating agents/drugs: α-CD3/CD28 beads (1:1 bead-to-cell ratio), TNF (10 ng/mL), 12□O□tetradecanoylphorbol□13□acetate at 10 μM concentration (TPA; Sigma□Aldrich, Saint Louis, MI, USA) and suberoyl anilide hydroxamic acid 0.5 μM (SAHA; Selleckchem, Houston, TX, USA S1047).

### Fluorescence lifetime imaging microscopy (FLIM)

Time-domain FLIM experiments on live TZM-bl cells expressing Laconic were performed using a Time Correlated Single Photon Counting (TCSPC) approach operated by the FALCON module (Leica Microsystems, Manheim, Germany) integrated within the Leica SP8-X-SMD microscope. Cells of interest were selected under a 63x/NA 1.20 water objective. Laconic-expressing cells were excited using a 440 nm pulsed laser, tuned at 40□MHz and subsequently detected by a hybrid internal detector (475nm − 510nm) in photon counting mode. Transfected cells co-expressing Laconic and Gag-mCherry were also excited using a DPSS 561 continuous laser, and mCherry fluorescence emission was detected by a hybrid internal detector (525nm − 560nm). Images were acquired using a scan speed of 200 Hz with a pixel size of 120 nm and obtained after 10 times frame repetitions.

### Image analysis

FLIM images were analyzed using the FALCON module (Leica Microsystems, Manheim, Germany) integrated within the Leica SP8-X-SMD microscope. Images were binned 3 x 3 to reach at least 100 counts/pixel. Individual cells expressing Laconic alone (No Env infected) or co-expressing Gag-mCherry were selected as regions of interest. A two-exponential decay deconvoluted with the Instrument Response Function (IRF) and fitted by a Marquandt nonlinear least-square algorithm was applied to the photon counting histogram with the long lifetime component fixed to 2.6 ns. The average lifetime, intensity weighted (TauInt) was calculated per cell and normalized to the mean TauInt of the no-Env condition of each experiment. Normalized results from three independent experiments were statistically analyzed using a one-way ANOVA test (OriginLab software, Northampton, USA).

### MTT assay of cell viability

Cell viability upon treatment with auranofin and/or BSO was measured using the CellTiter 96^®^Non□Radioactive Cell Proliferation Assay (MTT) (Promega; Madison, WI, USA) according to the manufacturer’s instructions, as described in (Shytaj *et al*, 2020). Absorbance values were measured using an Infinite 200 PRO (Tecan, Männedorf, Switzerland) plate reader. After blank subtraction, absorbance values were normalized using matched untreated controls.

### Flow cytometry and cell sorting

To measure GFP expression in J-Lat 9.2 cells, 500 × 10^5^ cells were fixed with 4% PFA in PBS, washed twice with PBS and resuspended in the FACS buffer. GFP fluorescence was measured using a BD FACSCelesta (Becton Dickinson, Franklin Lakes, NJ, USA) flow cytometer and analyzed using the FlowJo software (FlowJo LLC, Ashland, Oregon, USA v7.6.5).

GFP expression in hµglia/HIV HC69 and HIV-1 infected iPSC-derived microglia was measured using a LSRFortessa instrument for cell sorting, the FACSDiva software (Becton Dickinson, Franklin Lakes, NJ, USA) for data collection, and the WinList 3D software (Verity Software House, Topsham, ME, USA) for data analysis.

Viability of U937 and U1 cells was assessed by propidium iodide staining. Briefly, cells were suspended in PBS, and stained with 1.5 µM propidium iodide (PI) for 15 min in the dark. After washing twice with PBS, cells were analyzed on a flow cytometer using the phycoerythrin detector with 488 nm excitation and 575/26 nm emission on a FACSVerse Flow cytometer (Becton Dickinson, Franklin Lakes, NJ, USA). CD4^+^ T-cells of PLWH were sorted based on LIVE/DEAD Cell Viability staining (Thermo Fisher, Waltham, MA, USA) using a LSR Aria cell sorter and the FACSDiva software (Becton Dickinson, Franklin Lakes, NJ, USA) for data collection.

### Redox Potential Measurement

Intracellular redox potential measurements in U1 cells were done as described earlier (Bhaskar *et al*, 2015). Briefly, the ratio-metric response of cells expressing the Grx1-roGFP2 sensor was obtained by measuring excitation at 405 and 488 nm at a fixed emission (510/10 nm) using a FACS Verse Flow cytometer (Becton Dickinson, Franklin Lakes, NJ, USA).

### Real-time PCR and ALU-HIV PCR

HIV-1 expression (Gag-p24) in U1 cells was measured by qPCR as described previously (Bhaskar *et al*, 2015). Briefly, total cellular RNA was reverse transcribed to cDNA (iScriptTM cDNA synthesis kit, Bio-Rad, Hercules, CA, United States). Real time PCR (iQTM SYBR Green Supermix, Bio-Rad Hercules, CA, United States), was performed using the Bio-Rad C1000TM real time PCR system. Expression level of Gag was normalized to human β-actin. All experiments were done at least twice in triplicate.

To measure integrated HIV-1 DNA in live CD4^+^ T-cells of PLWH, Alu-HIV PCR was performed as described in (Chomont *et al*, 2009). HIV-1 reactivation in the same cell types was measured by *Tat/rev* Induced Limiting Dilution Assay as described in (Procopio *et al*, 2015).

### Statistical analysis

Statistical analysis of *in-silico* data is described in the respective Methods subchapters. HIV-1 reactivation and cell viability data were analyzed by parametric (*i*.*e*. one□ or two□way ANOVA tests) or non-parametric (*i*.*e*. Friedman test). Parametric testing was adopted when normality could be hypothesized (sample size ≤ 3) or restored through an appropriate transformation (*e*.*g*. tangent in the case of Figure 3B). Data sets characterized by sample size > 3 which did not pass the normality tests (D’Agostino & Pearson or Shapiro-Wilk) and for which a transformation was not applicable were analyzed by non-parametric tests. For both parametric and non-parametric tests, post-test comparisons were used to compare specific groups as described in the Figure captions. Analyses were performed using GraphPad Prism (GraphPad Software, San Diego, CA, USA).

## Acknowledgments

R.S.D. acknowledges support from the Fundação de Amparo à Pesquisa do Estado de São Paulo and the Conselho Nacional de Desenvolvimento Científico e Tecnológico (FAPESP 2013/11323-5; CNPq − 454700-2014-8; CNPq/DECIT 441817/2018-1). I.L.S. acknowledges support from the Humboldt Foundation (Ref 3.3 - ITA - 1193954 - HFST-P) and the Fundação de Amparo à Pesquisa do Estado de São Paulo (CNPJ 43.828.151/0001-45). A.Si. acknowledges support from the Department of Biotechnology, Indian Institute of Science (# 22-0905-0006-05-987-436). The authors thank the Microarray Unit of the Genomics and Proteomics Core Facility, German Cancer Research Center (DKFZ), for providing Expression Profiling services.

## Authors’ Contributions

I.L.S., D.AC., A.Sa. conceived the project. I.L.S, F.A.P., S.PP, A.Si., M.L. R.S.D, D.AC., A.Sa. designed the experiments. I.L.S, F.A.P., I.CA, M.H.M., N.C., B.L, D.AC., performed *in-vitro* experiments. I.L.S., F.A.P., I.CA., D.AC., A.Sa. analyzed *in-vitro* data. M.T., M.F., K.L, F.Y., S.B., analyzed transcriptomic data. M.T. HY.T., A.R.G., analyzed metabolomic data. I.L.S, A.Sa. wrote the manuscript.

## Additional Files

**Additional File 1. Microarray analysis of the effect of HIV-1 infection on gene expression in CD4**^**+**^ **T-cells**. Primary CD4^+^ T-cells were isolated from total blood of healthy individuals, activated for 72 h with α-CD3-CD28 beads and infected with HIV-1_pNL4-3_ or mock-infected. Both infected and mock-infected cells were cultured for two weeks to model different infection stages as described in (Shytaj *et al*, 2020). On days 7, 9 and 14 post-infection, HIV-1 infected and mock-infected cells were subjected to microarray analysis. Data from different time points were pooled for the analysis. Significantly enriched pathways in HIV-1 infected and mock-infected cells were identified by Gene Set Enrichment Analysis [GSEA (Subramanian *et al*, 2005)]. Number of donors = 2.

**Additional File 2. Modulation of glycolytic pathway in the CD4**^**+**^ **T-cell proteome**. Heatmap of the expression of the glycolytic pathway in HIV-1 infected vs mock-infected cells. Cells were activated for 72 h with α-CD3-CD28 beads and infected with HIV-1_pNL4-3_ or mock-infected. Both infected and mock-infected cells were cultured for two weeks to model different infection stages as described in (Shytaj *et al*, 2020). On days 3, 7, 9 and 14 post-infection, HIV-1 infected and mock-infected cells were subjected to proteomic analysis (Shytaj *et al*, 2020). Data were analyzed by Student’s t-test after pooling the different time points.

**Additional File 3. Modulation of glycolytic pathway transcription during productive or latent HIV-1 infection of microglia**. Heatmap of the relative gene expression of the glycolytic pathway in C20/HC69 microglial cells (Garcia-Mesa *et al*, 2017) is shown. Microglial cells were subjected to RNA-Seq analysis under four conditions: uninfected (C20), uninfected and activated (C20-TNF), latently HIV-1 infected (HC69) and reactivated HIV-1 infection (HC69-TNF). Expression data were analyzed by Deseq2 (Love *et al*, 2014).

**Additional File 4. Effect of the glycolysis depressing agent dexamethasone (DEXA) on the baseline HIV-1 expression in HC69 microglia cells**. HIV-1 infected microglia cells (HC69) were left untreated or treated with DEXA (1μM) for 72 h and then subjected to scRNA-Seq. The bubble plot depicts the percentage of cells in which the transcriptional expression of HIV-1 or the genes of the HUMAN-GLYCOLYSIS pathway was detectable.

**Additional Files 5 and 6. List of metabolic pathways enriched in latently (Additional file 5) and productively (Additional file 6) infected microglial cells**. Microglial cells were subjected to metabolomic analysis under four conditions: uninfected (C20), uninfected and activated (C20-TNF), latently HIV-1 infected (HC69) and reactivated HIV-1 infection (HC69-TNF). Each infected condition was compared to its uninfected counterpart. The analysis was performed using MetaboAnalyst (Xia et al. 2009).

**Additional File 7. Auranofin (AF) and buthionine sulfoximine (BSO) co-treatment induces synergistic oxidative stress in U1 cells**. U1 cells expressing the redox sensor Grx1-roGFP2 (Bhaskar *et al*, 2015) were exposed to various combinations of AF and/or BSO and the roGFP2 ratio (405/488 nm) was measured 24 h post-treatment. Fold change was calculated by dividing the roGFP2 ratios under treated conditions with the untreated cells. Data (mean ± SD of three experiments) were analyzed by One Way ANOVA followed by Tukey’s post-test (only comparisons between matching drug dosages are shown for simplicity’s sake).

* p< 0.05, **** p< 0.0001.

**Additional Files 8 and 9. HIV-1 reactivation (A) and viability (B) in lymphoid (Additional file 8) and myeloid (Additional file 9) cells following treatment with auranofin (AF) and buthionine sulfoximine (BSO)**. Culturing conditions for each cell type are detailed in the Materials and Methods section. Cells were left untreated or treated with AF (500 nM), BSO (250 μM), or a combination of the two. Treatment duration was 48 h except for Jurkat/J-lat 9.2, Th17 and U937/U1 cells that were treated for 24 h. HIV-1 expression was determined by FACS (Th17, J-Lat9.2, HC69 microglia, iGM microglia, THP-1) or qPCR (U1). Data are expressed as mean ± SD of three replicates and were analyzed by One-Way ANOVA followed by Tukey’s post-test (A) or Two-Way ANOVA followed by Sidak’s post-test (B).

* p< 0.05, ** p< 0.01, *** p< 0.001, **** p< 0.0001.

**Additional File 10. Modulation of glycolytic metabolites in cells of PLWH treated with auranofin (AF) and/or buthionine sulfoximine (BSO)**. Peripheral blood mononuclear cells (PBMC), isolated from total blood of PLWH, were treated with auranofin AF (500 nM), BSO (250 μM) or a combination of the two, for 24 h. PBMC donors were selected from PLWH enrolled in trial NCT02961829 (Diaz *et al*, 2019). Cells were subjected to metabolomic analysis and the top enriched pathways were ordered according to p values obtained with Q statistic for metabolic datasets performed with Globaltest (MetaboAnalyst) (Xia *et al*, 2009). Number of donors = 6.

**Additional File 11. HIV-1 reactivation in CD4**^**+**^ **T-cells of PLWH under ART following treatment with auranofin (AF) and buthionine sulfoximine (BSO)**. CD4^+^ T-cells were isolated from total blood of PLWH under suppressive ART and left untreated or treated with AF (500 nM), BSO (250 μM) or a combination of the two for 48 h. Viral reactivation was measured by Tat/rev Induced Limiting Dilution Assay [TILDA (CD4^+^ T-cells of PLWH)] (Procopio *et al*, 2015). Data are expressed as medians and interquartile ranges.

