## Supplementary figures and images for "Glycolysis downregulation is a hallmark of HIV-1 latency and sensitizes infected cells to oxidative stress"

### Additional File 2

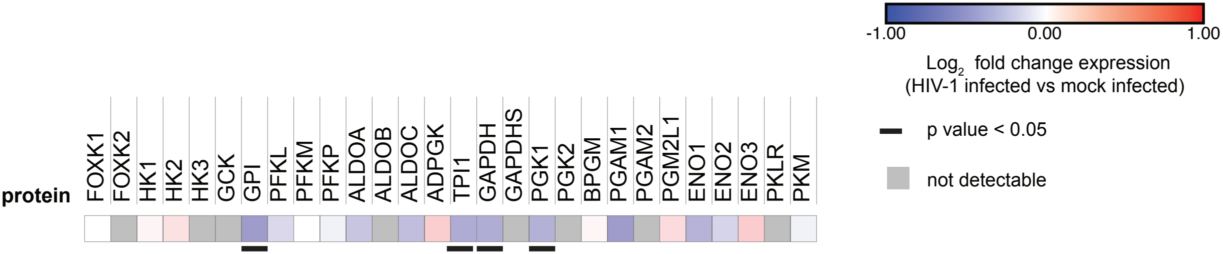

### Additional File 3

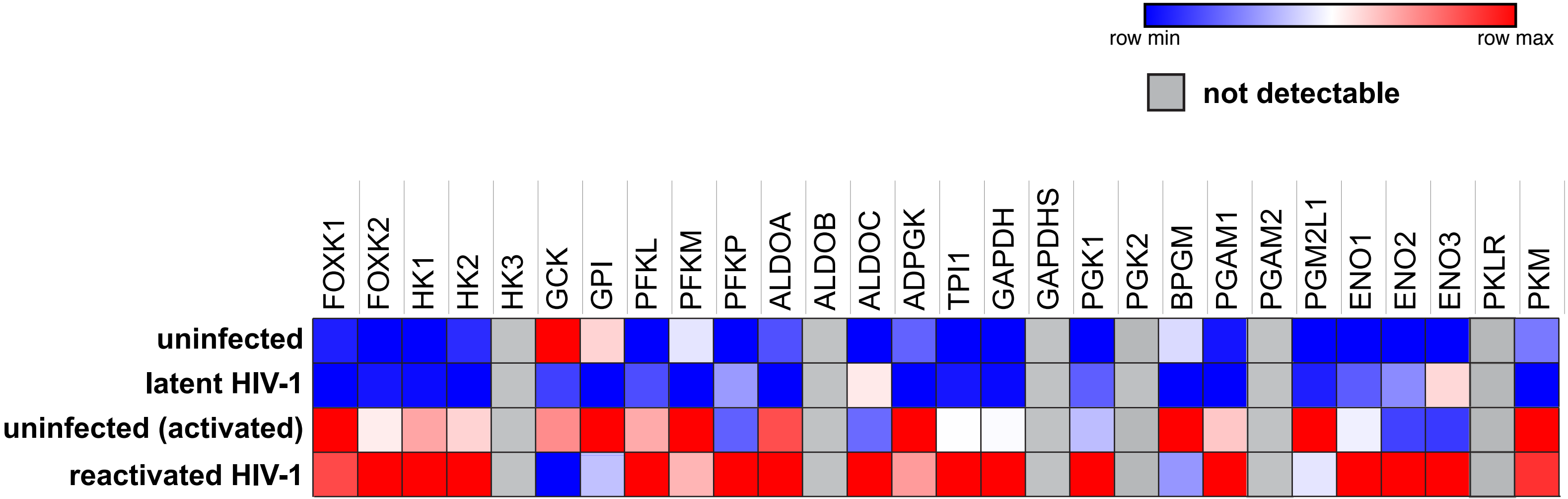

### Additional File 4

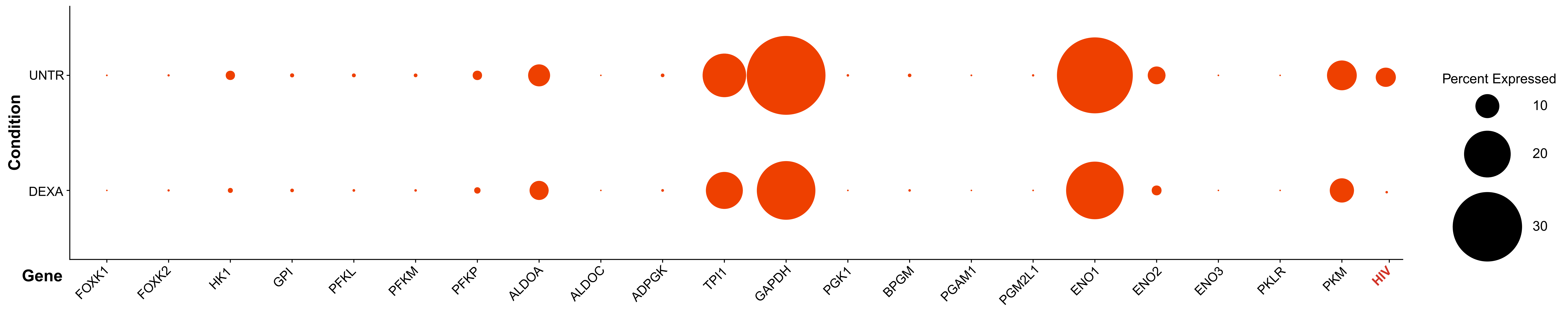

### Additional File 7

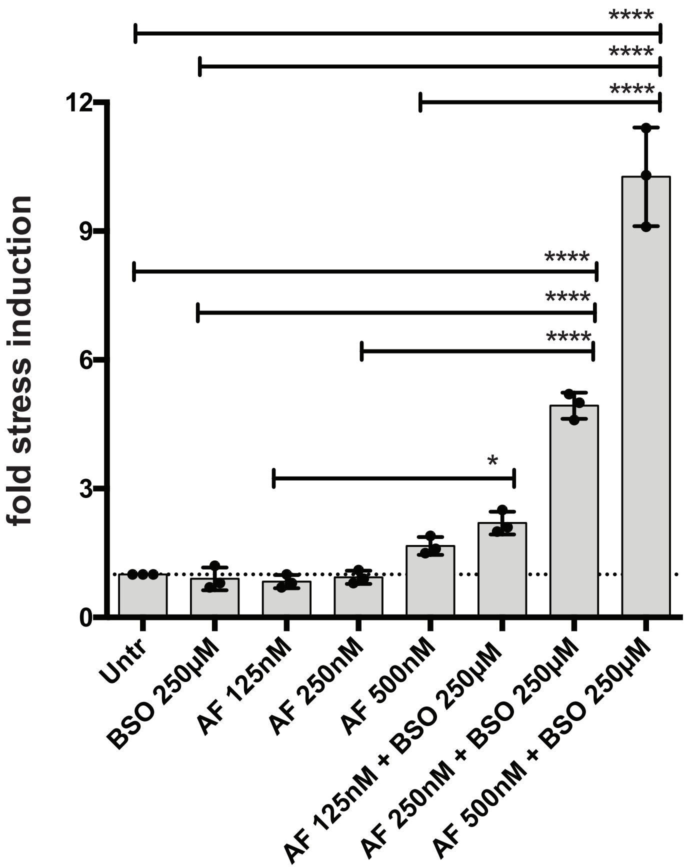

### Additional File 8

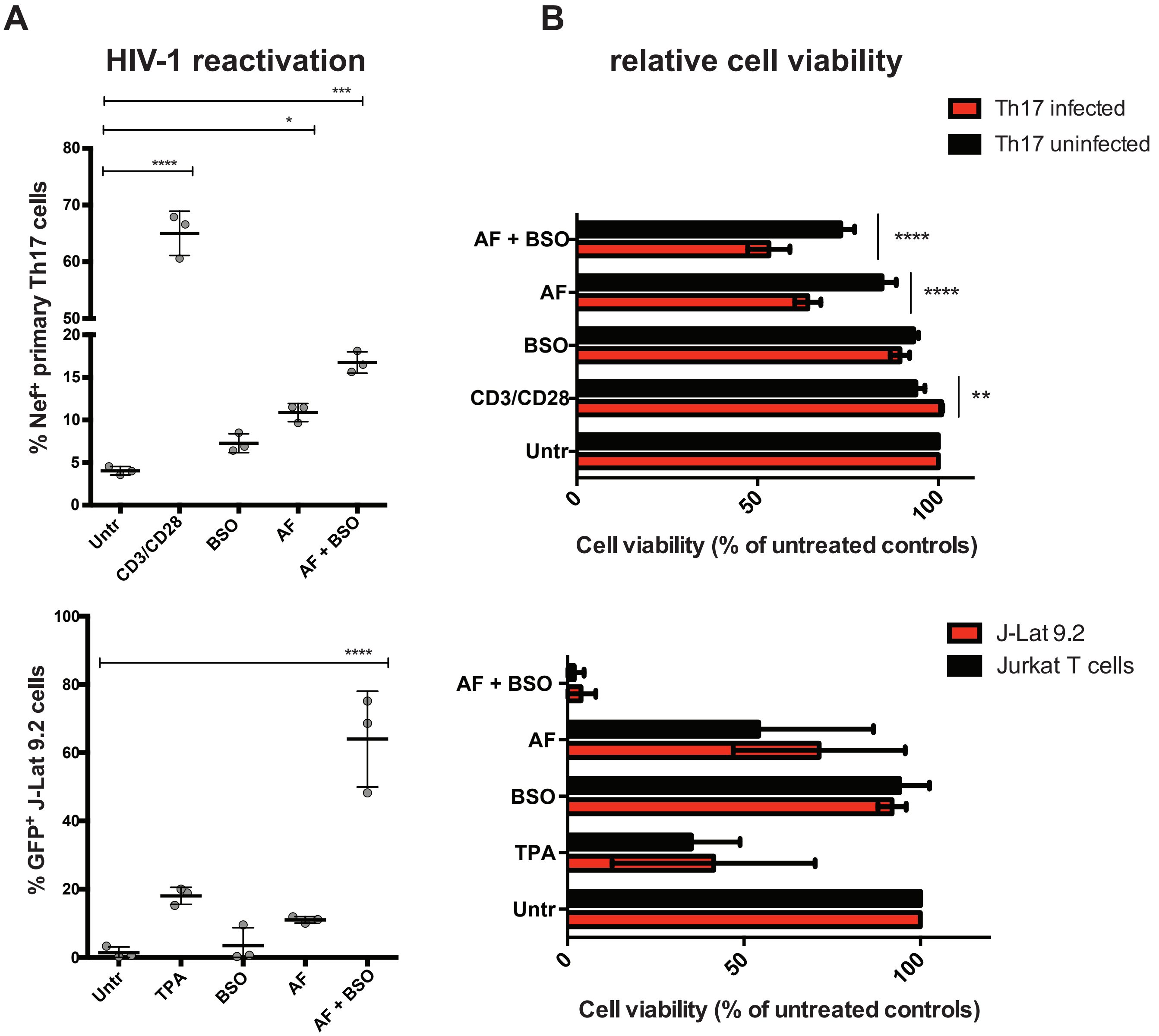

### Additional File 9

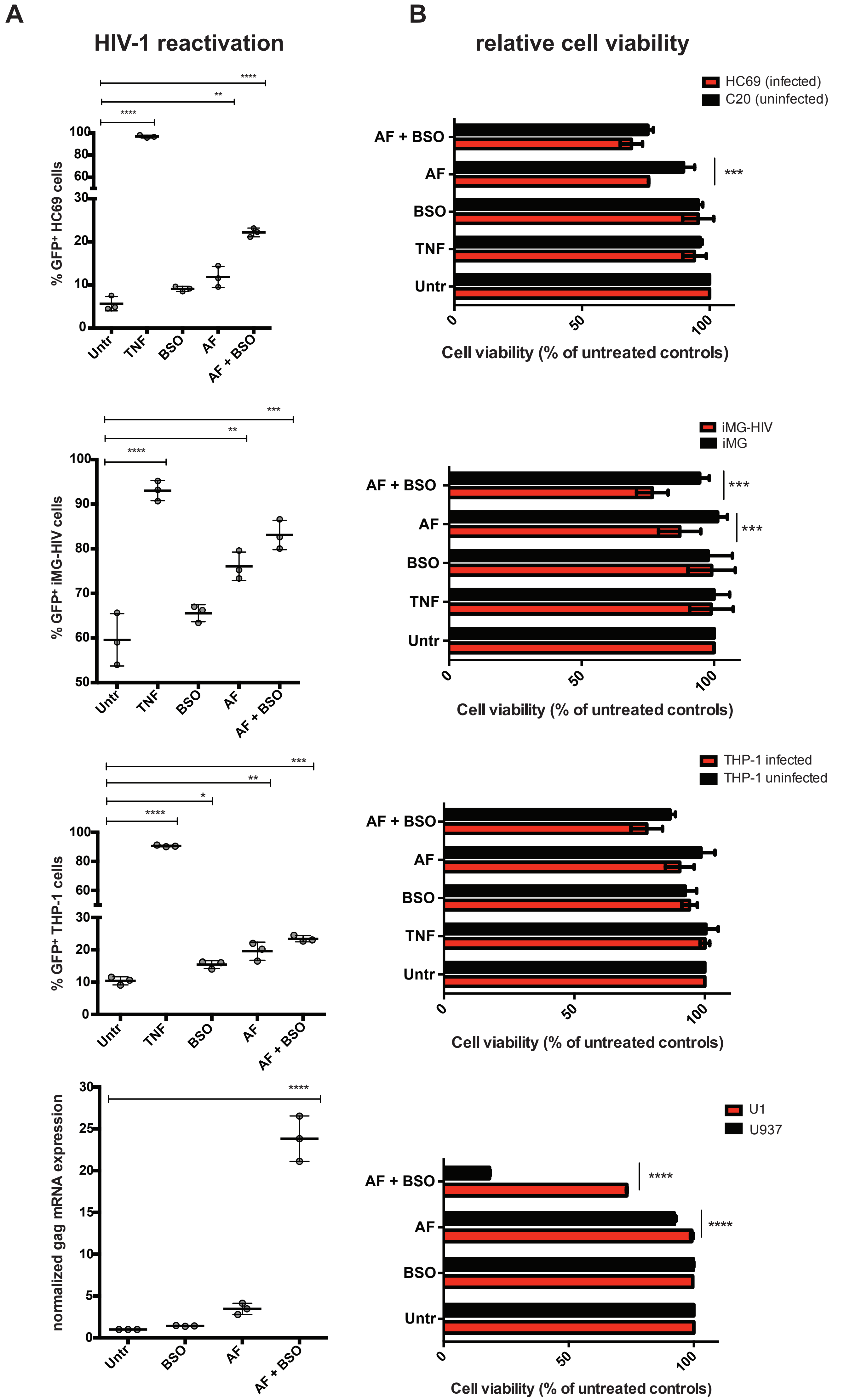

### Additional File 10

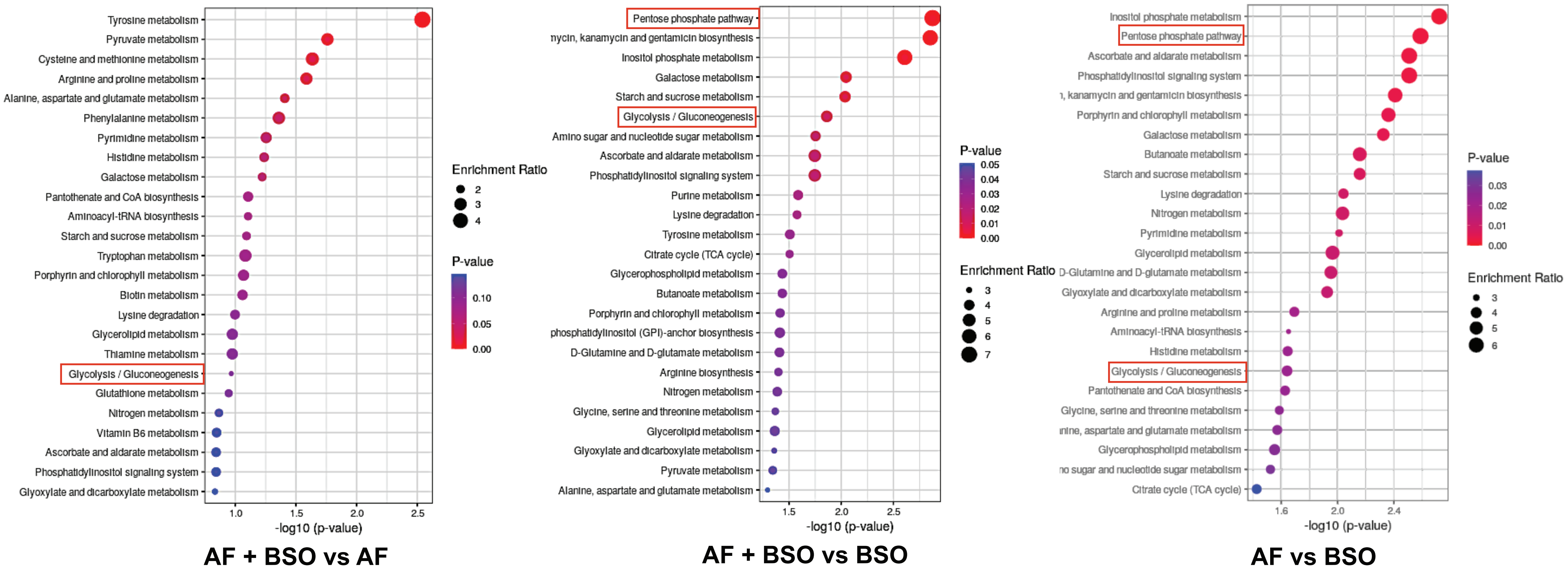

### Additional File 11

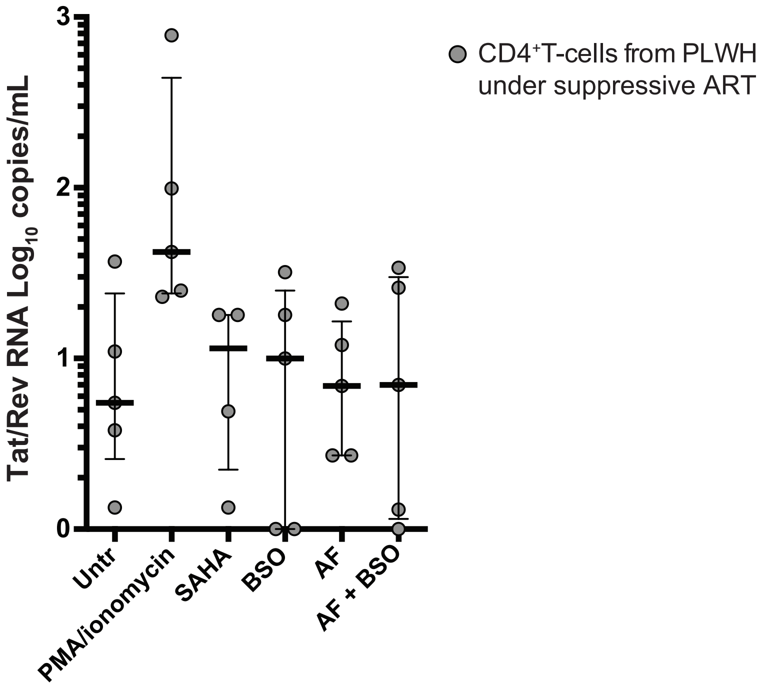
