## Additional File 5 for "Glycolysis downregulation is a hallmark of HIV-1 latency and sensitizes infected cells to oxidative stress"

### Quantitative Enrichment Analysis (latent infection)

| | Total Cmpd | Hits | Statistic Q | Expected Q | Raw p | Holm p | FDR |
|---|---|---|---|---|---|---|---|
| Pyrimidine metabolism | 39 | 11 | 97.96 | 20.00 | 4.54E-07 | 2.81E-05 | 2.81E-05 |
| Purine metabolism | 65 | 18 | 96.93 | 20.00 | 2.23E-06 | 1.36E-04 | 4.31E-05 |
| Glyoxylate and dicarboxylate metabolism | 32 | 8 | 96.29 | 20.00 | 2.75E-06 | 1.65E-04 | 4.31E-05 |
| Aminoacyl-tRNA biosynthesis | 48 | 17 | 93.68 | 20.00 | 2.78E-06 | 1.65E-04 | 4.31E-05 |
| Glycine, serine and threonine metabolism | 33 | 10 | 97.72 | 20.00 | 5.85E-06 | 3.39E-04 | 5.24E-05 |
| Glutathione metabolism | 28 | 8 | 90.47 | 20.00 | 6.85E-06 | 3.90E-04 | 5.24E-05 |
| Alanine, aspartate and glutamate metabolism | 28 | 12 | 95.74 | 20.00 | 7.27E-06 | 4.07E-04 | 5.24E-05 |
| Porphyrin and chlorophyll metabolism | 30 | 2 | 99.03 | 20.00 | 7.68E-06 | 4.22E-04 | 5.24E-05 |
| Sphingolipid metabolism | 21 | 1 | 99.54 | 20.00 | 7.84E-06 | 4.23E-04 | 5.24E-05 |
| Arginine biosynthesis | 14 | 10 | 94.69 | 20.00 | 8.60E-06 | 4.56E-04 | 5.24E-05 |
| Pyruvate metabolism | 22 | 5 | 85.77 | 20.00 | 9.30E-06 | 4.84E-04 | 5.24E-05 |
| Citrate cycle (TCA cycle) | 20 | 7 | 96.08 | 20.00 | 1.28E-05 | 6.51E-04 | 6.60E-05 |
| Primary bile acid biosynthesis | 46 | 2 | 99.05 | 20.00 | 1.59E-05 | 7.95E-04 | 7.17E-05 |
| Tyrosine metabolism | 42 | 4 | 91.19 | 20.00 | 1.62E-05 | 7.95E-04 | 7.17E-05 |
| Pentose phosphate pathway | 22 | 6 | 82.66 | 20.00 | 2.18E-05 | 1.05E-03 | 8.49E-05 |
| D-Glutamine and D-glutamate metabolism | 6 | 5 | 98.15 | 20.00 | 2.19E-05 | 1.05E-03 | 8.49E-05 |
| Butanoate metabolism | 15 | 5 | 95.85 | 20.00 | 2.80E-05 | 1.29E-03 | 1.02E-04 |
| Cysteine and methionine metabolism | 33 | 11 | 90.85 | 20.00 | 5.58E-05 | 2.51E-03 | 1.92E-04 |
| Arachidonic acid metabolism | 36 | 1 | 98.51 | 20.00 | 8.34E-05 | 3.67E-03 | 2.72E-04 |
| Taurine and hypotaurine metabolism | 8 | 4 | 82.63 | 20.00 | 1.11E-04 | 4.76E-03 | 3.44E-04 |
| Ascorbate and aldarate metabolism | 8 | 3 | 95.44 | 20.00 | 1.38E-04 | 5.80E-03 | 4.08E-04 |
| Glycolysis / Gluconeogenesis | 26 | 8 | 77.34 | 20.00 | 1.70E-04 | 6.96E-03 | 4.78E-04 |
| Arginine and proline metabolism | 38 | 10 | 79.92 | 20.00 | 1.94E-04 | 7.77E-03 | 5.23E-04 |
| Lysine degradation | 25 | 5 | 95.66 | 20.00 | 2.33E-04 | 9.09E-03 | 6.02E-04 |
| Biosynthesis of unsaturated fatty acids | 36 | 8 | 78.81 | 20.00 | 2.47E-04 | 9.39E-03 | 6.13E-04 |
| Galactose metabolism | 27 | 5 | 86.22 | 20.00 | 2.89E-04 | 1.07E-02 | 6.88E-04 |
| Biotin metabolism | 10 | 1 | 97.05 | 20.00 | 3.29E-04 | 1.18E-02 | 7.55E-04 |
| Inositol phosphate metabolism | 30 | 4 | 75.46 | 20.00 | 3.61E-04 | 1.27E-02 | 7.78E-04 |
| Histidine metabolism | 16 | 4 | 91.37 | 20.00 | 3.64E-04 | 1.27E-02 | 7.78E-04 |
| Amino sugar and nucleotide sugar metabolism | 37 | 6 | 87.46 | 20.00 | 3.81E-04 | 1.27E-02 | 7.87E-04 |
| Tryptophan metabolism | 41 | 6 | 47.38 | 20.00 | 9.47E-04 | 3.03E-02 | 1.89E-03 |
| Retinol metabolism | 17 | 1 | 93.05 | 20.00 | 1.85E-03 | 5.75E-02 | 3.59E-03 |
| Phosphatidylinositol signaling system | 28 | 1 | 90.29 | 20.00 | 3.66E-03 | 1.10E-01 | 6.87E-03 |
| beta-Alanine metabolism | 21 | 4 | 55.44 | 20.00 | 4.51E-03 | 1.31E-01 | 8.20E-03 |
| Fructose and mannose metabolism | 20 | 2 | 83.03 | 20.00 | 4.70E-03 | 1.32E-01 | 8.20E-03 |
| Valine, leucine and isoleucine biosynthesis | 8 | 3 | 83.18 | 20.00 | 4.76E-03 | 1.32E-01 | 8.20E-03 |
| Neomycin, kanamycin and gentamicin biosynthesis | 2 | 1 | 87.47 | 20.00 | 6.15E-03 | 1.60E-01 | 1.03E-02 |
| Pentose and glucuronate interconversions | 18 | 2 | 69.90 | 20.00 | 8.01E-03 | 2.00E-01 | 1.30E-02 |
| Nicotinate and nicotinamide metabolism | 15 | 4 | 52.46 | 20.00 | 8.15E-03 | 2.00E-01 | 1.30E-02 |
| Glycerophospholipid metabolism | 36 | 4 | 80.58 | 20.00 | 8.36E-03 | 2.00E-01 | 1.30E-02 |
| Starch and sucrose metabolism | 18 | 2 | 74.78 | 20.00 | 1.24E-02 | 2.72E-01 | 1.87E-02 |
| Valine, leucine and isoleucine degradation | 40 | 3 | 60.30 | 20.00 | 1.46E-02 | 3.06E-01 | 2.15E-02 |
| Glycerolipid metabolism | 16 | 1 | 79.05 | 20.00 | 1.78E-02 | 3.55E-01 | 2.56E-02 |
| Pantothenate and CoA biosynthesis | 19 | 5 | 58.11 | 20.00 | 1.83E-02 | 3.55E-01 | 2.59E-02 |
| Glycosylphosphatidylinositol (GPI)-anchor biosynthesis | 14 | 1 | 76.68 | 20.00 | 2.22E-02 | 4.00E-01 | 3.06E-02 |
| Phenylalanine metabolism | 10 | 2 | 49.22 | 20.00 | 5.67E-02 | 9.64E-01 | 7.48E-02 |
| Phenylalanine, tyrosine and tryptophan biosynthesis | 4 | 2 | 49.22 | 20.00 | 5.67E-02 | 9.64E-01 | 7.48E-02 |
| Linoleic acid metabolism | 5 | 1 | 52.08 | 20.00 | 1.05E-01 | 1.00E+00 | 1.36E-01 |
| Selenocompound metabolism | 20 | 1 | 38.34 | 20.00 | 1.90E-01 | 1.00E+00 | 2.40E-01 |
| Fatty acid degradation | 39 | 3 | 29.02 | 20.00 | 1.99E-01 | 1.00E+00 | 2.47E-01 |
| Folate biosynthesis | 27 | 1 | 33.95 | 20.00 | 2.25E-01 | 1.00E+00 | 2.73E-01 |
| alpha-Linolenic acid metabolism | 13 | 1 | 31.26 | 20.00 | 2.49E-01 | 1.00E+00 | 2.97E-01 |
| Vitamin B6 metabolism | 9 | 1 | 22.81 | 20.00 | 3.38E-01 | 1.00E+00 | 3.96E-01 |
| Ubiquinone and other terpenoid-quinone biosynthesis | 9 | 1 | 18.36 | 20.00 | 3.97E-01 | 1.00E+00 | 4.55E-01 |
| Riboflavin metabolism | 4 | 1 | 10.56 | 20.00 | 5.30E-01 | 1.00E+00 | 5.97E-01 |
| Nitrogen metabolism | 6 | 2 | 12.76 | 20.00 | 5.99E-01 | 1.00E+00 | 6.64E-01 |
| Thiamine metabolism | 7 | 1 | 5.21 | 20.00 | 6.63E-01 | 1.00E+00 | 7.22E-01 |
| Synthesis and degradation of ketone bodies | 5 | 1 | 3.53 | 20.00 | 7.21E-01 | 1.00E+00 | 7.45E-01 |
| Propanoate metabolism | 23 | 1 | 3.53 | 20.00 | 7.21E-01 | 1.00E+00 | 7.45E-01 |
| Terpenoid backbone biosynthesis | 18 | 1 | 3.53 | 20.00 | 7.21E-01 | 1.00E+00 | 7.45E-01 |
| Fatty acid elongation | 39 | 2 | 1.51 | 20.00 | 9.49E-01 | 1.00E+00 | 9.64E-01 |
| Fatty acid biosynthesis | 47 | 4 | 1.03 | 20.00 | 9.98E-01 | 1.00E+00 | 9.98E-01 |
