## Additional File 6 for "Glycolysis downregulation is a hallmark of HIV-1 latency and sensitizes infected cells to oxidative stress"

### Quantitative Enrichment Analysis (productive infection)

| | Total Cmpd | Hits | Statistic Q | Expected Q | Raw p | Holm p | FDR |
|---|---|---|---|---|---|---|---|
| Pyrimidine metabolism | 39 | 11 | 98.41 | 20.00 | 5.91E-07 | 3.67E-05 | 1.60E-05 |
| Alanine, aspartate and glutamate metabolism | 28 | 12 | 94.09 | 20.00 | 7.28E-07 | 4.44E-05 | 1.60E-05 |
| Purine metabolism | 65 | 18 | 95.00 | 20.00 | 8.24E-07 | 4.94E-05 | 1.60E-05 |
| Ascorbate and aldarate metabolism | 8 | 3 | 99.20 | 20.00 | 1.03E-06 | 6.09E-05 | 1.60E-05 |
| Citrate cycle (TCA cycle) | 20 | 7 | 97.20 | 20.00 | 3.06E-06 | 1.78E-04 | 2.49E-05 |
| Tyrosine metabolism | 42 | 4 | 93.95 | 20.00 | 3.09E-06 | 1.78E-04 | 2.49E-05 |
| Glycolysis / Gluconeogenesis | 26 | 8 | 88.58 | 20.00 | 3.13E-06 | 1.78E-04 | 2.49E-05 |
| Inositol phosphate metabolism | 30 | 4 | 97.55 | 20.00 | 3.35E-06 | 1.84E-04 | 2.49E-05 |
| Butanoate metabolism | 15 | 5 | 95.18 | 20.00 | 3.61E-06 | 1.95E-04 | 2.49E-05 |
| D-Glutamine and D-glutamate metabolism | 6 | 5 | 96.93 | 20.00 | 7.25E-06 | 3.84E-04 | 3.94E-05 |
| Cysteine and methionine metabolism | 33 | 11 | 94.81 | 20.00 | 7.35E-06 | 3.84E-04 | 3.94E-05 |
| Arginine biosynthesis | 14 | 10 | 90.83 | 20.00 | 7.63E-06 | 3.89E-04 | 3.94E-05 |
| Fructose and mannose metabolism | 20 | 2 | 92.56 | 20.00 | 1.00E-05 | 5.01E-04 | 4.78E-05 |
| Glycine, serine and threonine metabolism | 33 | 10 | 97.37 | 20.00 | 1.15E-05 | 5.65E-04 | 5.10E-05 |
| Lysine degradation | 25 | 5 | 97.79 | 20.00 | 1.28E-05 | 6.14E-04 | 5.29E-05 |
| Pyruvate metabolism | 22 | 5 | 83.79 | 20.00 | 1.61E-05 | 7.58E-04 | 6.25E-05 |
| Amino sugar and nucleotide sugar metabolism | 37 | 6 | 88.40 | 20.00 | 1.92E-05 | 8.81E-04 | 6.98E-05 |
| Aminoacyl-tRNA biosynthesis | 48 | 17 | 80.97 | 20.00 | 3.39E-05 | 1.52E-03 | 1.13E-04 |
| Pentose phosphate pathway | 22 | 6 | 96.38 | 20.00 | 3.45E-05 | 1.52E-03 | 1.13E-04 |
| Galactose metabolism | 27 | 5 | 87.17 | 20.00 | 6.24E-05 | 2.69E-03 | 1.89E-04 |
| Phosphatidylinositol signaling system | 28 | 1 | 98.69 | 20.00 | 6.46E-05 | 2.71E-03 | 1.89E-04 |
| Glyoxylate and dicarboxylate metabolism | 32 | 9 | 86.50 | 20.00 | 6.74E-05 | 2.76E-03 | 1.89E-04 |
| Ubiquinone and other terpenoid-quinone biosynthesis | 9 | 1 | 98.64 | 20.00 | 7.01E-05 | 2.81E-03 | 1.89E-04 |
| Pentose and glucuronate interconversions | 18 | 2 | 93.24 | 20.00 | 1.03E-04 | 4.02E-03 | 2.66E-04 |
| Sphingolipid metabolism | 21 | 1 | 97.43 | 20.00 | 2.50E-04 | 9.48E-03 | 6.19E-04 |
| Starch and sucrose metabolism | 18 | 2 | 86.90 | 20.00 | 2.62E-04 | 9.68E-03 | 6.24E-04 |
| Histidine metabolism | 16 | 4 | 87.23 | 20.00 | 2.77E-04 | 9.99E-03 | 6.37E-04 |
| Glycerophospholipid metabolism | 36 | 4 | 81.29 | 20.00 | 4.00E-04 | 1.40E-02 | 8.86E-04 |
| Tryptophan metabolism | 41 | 6 | 86.39 | 20.00 | 4.61E-04 | 1.57E-02 | 9.85E-04 |
| Neomycin, kanamycin and gentamicin biosynthesis | 2 | 1 | 95.60 | 20.00 | 7.36E-04 | 2.43E-02 | 1.52E-03 |
| beta-Alanine metabolism | 21 | 4 | 79.05 | 20.00 | 8.10E-04 | 2.59E-02 | 1.62E-03 |
| Arginine and proline metabolism | 38 | 10 | 63.62 | 20.00 | 9.61E-04 | 2.98E-02 | 1.86E-03 |
| Taurine and hypotaurine metabolism | 8 | 4 | 87.82 | 20.00 | 1.04E-03 | 3.12E-02 | 1.96E-03 |
| Valine, leucine and isoleucine biosynthesis | 8 | 3 | 89.56 | 20.00 | 1.98E-03 | 5.73E-02 | 3.61E-03 |
| Primary bile acid biosynthesis | 46 | 2 | 90.98 | 20.00 | 2.51E-03 | 7.02E-02 | 4.44E-03 |
| Valine, leucine and isoleucine degradation | 40 | 3 | 77.81 | 20.00 | 3.81E-03 | 1.03E-01 | 6.57E-03 |
| Porphyrin and chlorophyll metabolism | 30 | 2 | 84.51 | 20.00 | 4.57E-03 | 1.19E-01 | 7.65E-03 |
| Pantothenate and CoA biosynthesis | 19 | 5 | 73.62 | 20.00 | 5.11E-03 | 1.28E-01 | 8.34E-03 |
| Nicotinate and nicotinamide metabolism | 15 | 4 | 79.44 | 20.00 | 5.65E-03 | 1.36E-01 | 8.99E-03 |
| Biosynthesis of unsaturated fatty acids | 36 | 8 | 58.00 | 20.00 | 6.60E-03 | 1.52E-01 | 1.02E-02 |
| Retinol metabolism | 17 | 1 | 86.83 | 20.00 | 6.81E-03 | 1.52E-01 | 1.03E-02 |
| Phenylalanine metabolism | 10 | 2 | 78.66 | 20.00 | 9.39E-03 | 1.97E-01 | 1.35E-02 |
| Phenylalanine, tyrosine and tryptophan biosynthesis | 4 | 2 | 78.66 | 20.00 | 9.39E-03 | 1.97E-01 | 1.35E-02 |
| Linoleic acid metabolism | 5 | 1 | 84.20 | 20.00 | 9.90E-03 | 1.97E-01 | 1.40E-02 |
| Glycosylphosphatidylinositol (GPI)-anchor biosynthesis | 14 | 1 | 80.67 | 20.00 | 1.50E-02 | 2.71E-01 | 2.07E-02 |
| Arachidonic acid metabolism | 36 | 1 | 71.36 | 20.00 | 3.43E-02 | 5.83E-01 | 4.62E-02 |
| Glutathione metabolism | 28 | 8 | 50.77 | 20.00 | 3.76E-02 | 6.01E-01 | 4.96E-02 |
| Folate biosynthesis | 27 | 1 | 58.99 | 20.00 | 7.44E-02 | 1.00E+00 | 9.62E-02 |
| Synthesis and degradation of ketone bodies | 5 | 1 | 53.42 | 20.00 | 9.89E-02 | 1.00E+00 | 1.20E-01 |
| Propanoate metabolism | 23 | 1 | 53.42 | 20.00 | 9.89E-02 | 1.00E+00 | 1.20E-01 |
| Terpenoid backbone biosynthesis | 18 | 1 | 53.42 | 20.00 | 9.89E-02 | 1.00E+00 | 1.20E-01 |
| Fatty acid degradation | 39 | 3 | 48.29 | 20.00 | 1.06E-01 | 1.00E+00 | 1.26E-01 |
| Nitrogen metabolism | 6 | 2 | 47.73 | 20.00 | 1.18E-01 | 1.00E+00 | 1.38E-01 |
| Thiamine metabolism | 7 | 1 | 44.63 | 20.00 | 1.47E-01 | 1.00E+00 | 1.69E-01 |
| Glycerolipid metabolism | 16 | 1 | 41.89 | 20.00 | 1.65E-01 | 1.00E+00 | 1.86E-01 |
| Selenocompound metabolism | 20 | 1 | 38.88 | 20.00 | 1.86E-01 | 1.00E+00 | 2.06E-01 |
| Fatty acid biosynthesis | 47 | 4 | 31.23 | 20.00 | 2.19E-01 | 1.00E+00 | 2.38E-01 |
| Riboflavin metabolism | 4 | 1 | 30.09 | 20.00 | 2.60E-01 | 1.00E+00 | 2.78E-01 |
| Fatty acid elongation | 39 | 2 | 27.51 | 20.00 | 2.82E-01 | 1.00E+00 | 2.97E-01 |
| alpha-Linolenic acid metabolism | 13 | 1 | 24.18 | 20.00 | 3.22E-01 | 1.00E+00 | 3.33E-01 |
| Biotin metabolism | 10 | 1 | 23.61 | 20.00 | 3.29E-01 | 1.00E+00 | 3.34E-01 |
| Vitamin B6 metabolism | 9 | 1 | 11.19 | 20.00 | 5.17E-01 | 1.00E+00 | 5.17E-01 |
